# Unifying the Research Landscape of Desiccation Tolerance to Identify Trends, Gaps, and Opportunities

**DOI:** 10.1101/2024.06.06.597802

**Authors:** Serena G. Lotreck, Mohammad Ghassemi, Robert T. VanBuren

## Abstract

Desiccation tolerance, or the ability to survive extreme dehydration, has evolved recurrently across the tree of life. While our understanding of the mechanisms underlying desiccation tolerance continues to expand, the compartmentalization of findings by study system impedes progress. Here, we analyzed 5,963 papers related to desiccation and examined model systems, research topics, citation networks, and disciplinary siloing over time. Our results show significant siloing, with plant science dominating the field, and relatively isolated clustering of plants, animal, microbial, and fungal literature. Topic modeling identified 46 distinct research topics, highlighting both commonalities and divergences across the knowledge of desiccation tolerance in different systems. We observed a rich diversity of model desiccation tolerant species within the community, contrasting the single species model for most biology research areas. To address citation gaps, we developed a rule-based algorithm to recommend new invitees to a niche conference, DesWorks, enhancing the integration of diverse research areas. The algorithm, which considers co-citation, co-authorship, research topics, and geographic data, successfully identified candidates with novel expertise that was unrepresented in previous conferences. Our findings underscore the importance of interdisciplinary collaboration in advancing desiccation tolerance research and provide a framework for using bibliometric tools to foster scientific integration.

## Introduction

Desiccation tolerance (DT) is the ability of an organism to survive near-complete dehydration (losing more than 95% of its cellular water content) for extended periods of time and to recover upon the re-addition of water (Morse et al., 2011). DT is found in all kingdoms of life (Alpert, 2005), including the seeds and pollen of flowering plants and vegetative tissues of diverse resurrection plants (Gaff and Oliver, 2013), gram-positive bacteria (Grzyb and Skłodowska, 2022), micro animals such as tardigrades, nematodes, rotifers, and artemia (Hibshman et al., 2020), and various lichen symbionts, yeast, and other fungi (Kranner et al., 2008). DT challenges the idea that water is required for all life at all times, and reveals unique mechanisms by which organisms can pause their metabolism in order to persist through long periods of dryness (Hoekstra et al., 2001). Additionally, given the pressures that a rapidly changing climate exerts on agriculture, stress biology, and particularly DT, is of great interest due to its potential applications (Koshland and Tapia, 2019).

Despite the phylogenetic breadth of organisms exhibiting DT, diverse lineages use similar biochemical and molecular mechanisms to survive life without water. These mechanisms include chaperones like late embryogenesis abundant (LEA) and heat shock proteins (HSPs), non-reducing osmoprotectants like trehalose and sucrose, and robust antioxidant systems. Many discoveries in the field were initially made in one system and subsequently found in others. For example, LEA proteins were first discovered in cotton seeds (Galau et al., 1986) and were later described in nematodes (Solomon et al., 2000; Browne et al., 2002), as well as numerous other plants, micro animals, fungi, protists, bacteria, and archaea (Battaglia et al., 2008; Boothby and Pielak, 2017). Similarly, the central role of trehalose in desiccation tolerance was discovered in micro animals and fungi, (Clegg and Filosa, 1961; Sussman, 1961; Clegg, 1965) and trehalose has been implicated for anhydrobiosis across multiple kingdoms. Despite these early cross kingdom discoveries, the field of desiccation tolerance has become increasingly compartmentalized as it grows, with research findings often confined to study systems or biological disciplines. This siloing is a problem in many scientific disciplines, but there is markedly little quantitative evidence supporting its prevalence or broader impact. Bibliometric analysis and natural language processing can quantify knowledge gaps across study systems, identify overlooked biological connections, predict future scientific collaborations and funding, and identify emerging research frontiers within the field (Wu et al., 2018; Yan et al., 2018; Makarov and Gerasimova, 2019; Resce et al., 2022).

Conferences play a crucial role in developing scientific connections and overcoming disciplinary silos. Papers presented at conferences are more likely to be cited by attendees, whether attendance is planned or incidental (Kim, Baek and Song, 2018). Conferences also create dedicated spaces for collaboration, enhancing the success of joint venues (Werker, Ooms, and Caniëls, 2016). However, collaborations are largely driven by the interpersonal social networks of scientists, as well as other factors like geographical proximity and economic similarity (Ma et al., 2014; Fernández et al., 2016; Werker et al., 2016). In chemistry, one study showed that corresponding authors knew at least one author from 75% of about 1,400 citations across 32 papers (Milard, 2014). The DT and anhydrobiosis research communities have few cross-disciplinary conferences, but the Desiccation Workshop (DesWorks) held every four years in South Africa since 1994, provides a key forum. Early DesWorks meetings were focused on DT in seeds, but the conference has since grown to include other organisms. Despite this growth, significant research gaps remain in biological disciplines and study systems, hindering the integration of important findings. Here, we propose that by increasing interpersonal connections across the field of DT, better integration in both citations and collaborations can be achieved.

This study aims to address three main questions. First, what is the extent of disciplinary siloing in DT research? To answer this question, we construct a citation network of 5,963 DT research papers from across the kingdoms of life, and classify them based on their study organism. Second, what topics are being studied in the DT literature, and how have these evolved over time? We use BERTopic, an unsupervised approach to define topic clusters in a body of literature, to examine the topics in DT research. Third, can we move beyond descriptive analysis and use bibliometric tools to help actively integrate the sub-disciplines within DT research? We developed a bibliometric-based algorithm to recommend new attendees to the DesWorks conference to fill research voids, utilizing co-citation, co-authorship, and topic modeling. By automating this process, we aim to overcome the challenge of information overload and enhance the integration and diversity within this specialized field. Our findings demonstrate that including our proposed attendees significantly improves the network integration of conference participants, advancing the overall field of DT research.

## Results

### Cataloging and exploring the desiccation tolerance literature

To explore the field of desiccation tolerance, we created a collection of desiccation tolerance literature (i.e., corpus) from the Web of Science. The broad search terms ‘desiccation tolerance’ and ‘anhydrobiosis’ returned 7,312 search results. Preprocessing and filtering to remove non-English results and articles related to soil science, materials science, and engineering, resulted in a final corpus of 5,963 papers on desiccation tolerance. Publication rate follows a rapid growth pattern that is characteristic of the scientific literature at large (González-Márquez et al., 2023), where most of the literature on desiccation tolerance was published over the last 20 years (Figure 1a).

**Figure 1.**
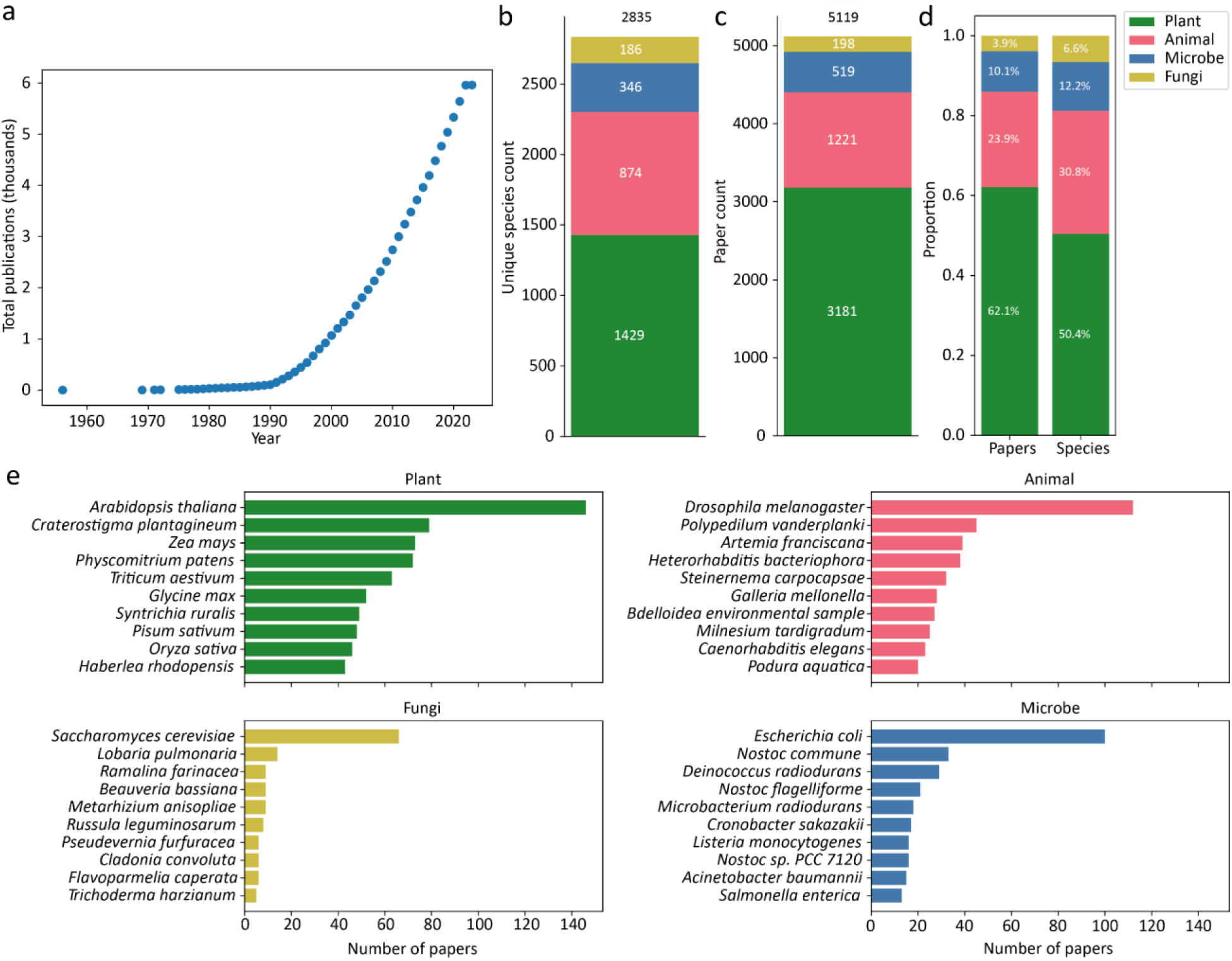
Overview of the desiccation tolerance literature. (a) Cumulative publications per year for publications related to “desiccation tolerance” OR “anhydrobiosis”. Note that 2023 is an incomplete year in the Web of Science XML Dataset, and total publications artificially levels off (b) Distribution of the desiccation tolerance literature for animal, plant, microbe, and fungi broken down by unique species count (b), total number of papers (c), and the proportion of both (d) as determined by TaxoNERD. (e) The top ten species with the most mentions in the corpus are shown for plant, animal, fungi, and microbe papers.

We first classified these papers based on study organisms using TaxoNERD with four broad categories: Plant (Plantae), Animal (Animalia), Microbe (Bacteria, Archaea, and Protozoa), and Fungi (Fungi). Of the 5,119 papers with abstracts that could be classified into study system groups, 3,181 belong to plant science, 1,221 focus on animals, 519 are microbial papers, and 198 study fungi (Figure 1c). Overall, 4,035 unique species can be found within the desiccation tolerance literature including 1,916 plants, 1,393 animals, 493 microbes, and 233 fungi (Figure 1b). While the majority of the research field is dominated by plants, the animal and fungi literature represent an outsized proportion of unique species compared to their relative representation in the dataset on a per-paper basis (Figure 1d).

Plant desiccation tolerance literature is dominated by work related to seed biology in the model species *Arabidopsis thaliana* and important grain crops and maize, wheat, soybean, rice, and pea (Figure 1e). *Craterostigma plantagineum* was established as the model resurrection plant in the 1990s (Bartels et al., 1990) and it is the most studied desiccation tolerant angiosperm. Other well-represented model resurrection plants include the bryophyte *Syntricia ruralis* and eudicot *Haberlea rhodopensis*. In animals, the most studied species include model anhydrobiotic animals such as tardigrades (*Milnesium tardigradum*), nematodes (*Heterorhabditis bateriophora, Steinernema carpocapsae*), midges (*Polypedilum vanderplanki*), rotifers (*Bdelloidea sp*.), and brine shrimp (*Artemia franiscana*). Yeast dominate the desiccation tolerant fungi literature followed by epiphytic lichen (*Lobaria pulmonaria, Ramalina farinacea*) and the insect-pathogenic fungus *Beauveria bassiana*. Aquatic and soil crust forming cyanobacteria and the model bacteria *E. coli* represent most of the microbe literature.

Paper keywords reflect disciplinary differences

Author-defined keywords are a simple way to categorize literature, and we explored the abundance of keywords in different disciplines and study systems and how they have changed over time. When we plot the absolute frequency of keywords over time (Figure 2a), we see that only “desiccation” and “desiccation tolerance” show a strong increase in usage, with most other keywords increasing slightly or staying about the same. However, normalizing keyword frequency by the number of papers published each year (Figure 2b) reveals that the use of “desiccation” and “desiccation tolerance”, in addition to all other keywords, has actually decreased over time. This can be explained by the introduction of new keywords as more papers are published. As more unique keywords are used, a relatively smaller share of total keyword usage is accounted for by each individual keyword. We see that approximately two new keywords have been introduced per published paper in the last 25 years, and that the overall number of new keywords each year continues to increase (Figure 2c). In general, keywords are becoming more specific as the field and knowledge of desiccation tolerance expand.

**Figure 2.**
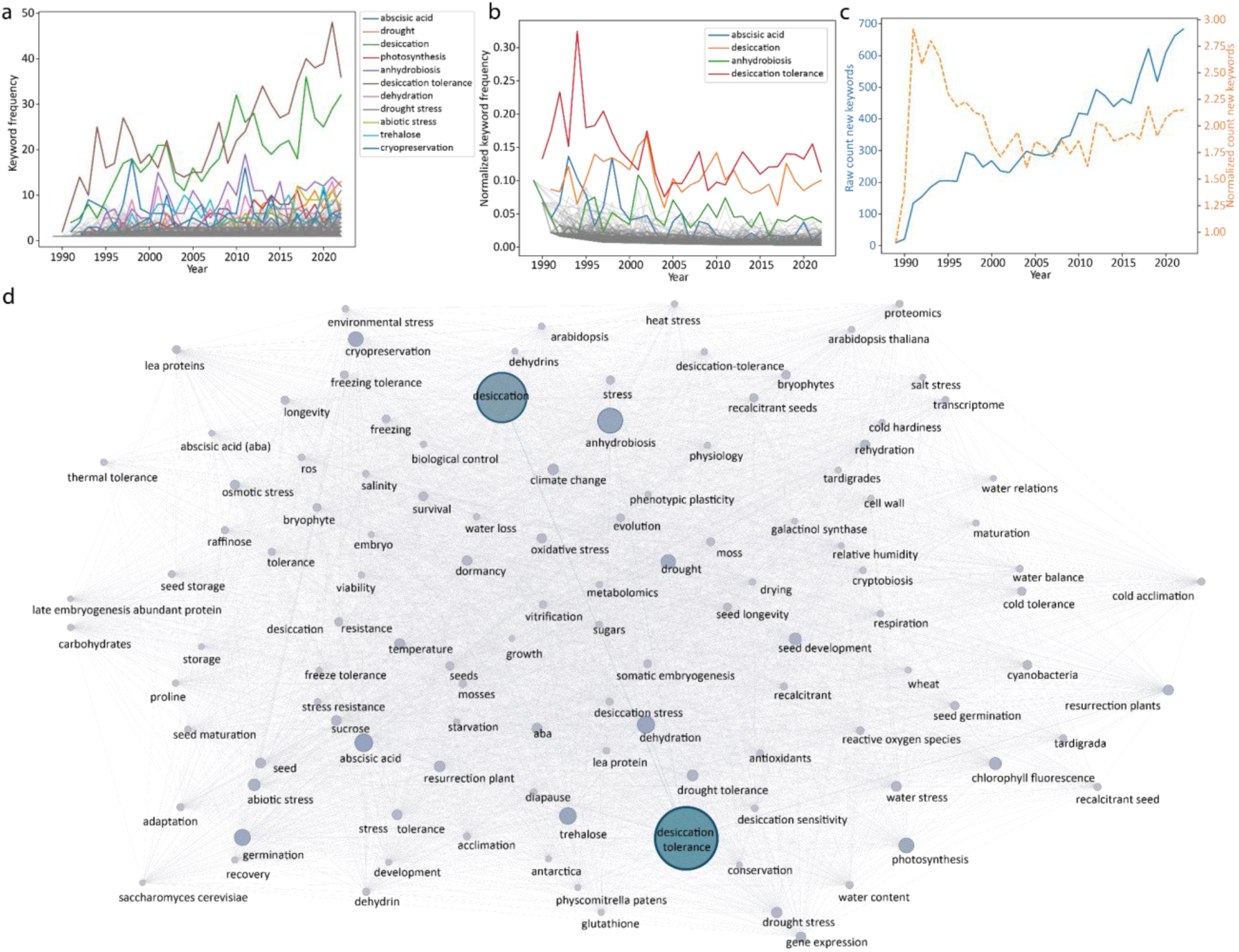
Dynamic use of keywords for the desiccation tolerance literature. (a) The absolute frequency of the top 1% most commonly used keywords is plotted over time, and the ten most abundant keywords are labeled individually. (b) Keyword frequency normalized by the number of papers per year, (c) The raw (blue, solid line) and normalized (orange, dotted line) number of new keywords introduced per year. Keywords with lower frequencies are grayed out for clarity in (a) and (b). (d) Keyword co-occurrence network for the top 1% of keywords. Nodes represent unique keywords and edges indicate that two keywords appeared together on the same paper, and are weighted by how frequently they co-occurred. The size of the nodes indicates the number of other nodes they are connected to (i.e., degree).

The keywords “desiccation” and “desiccation tolerance” are the only top 10 keywords shared across all four of our defined species groups (Table 1). Interestingly, the keywords “trehalose” and “anhydrobiosis” are used frequently in all groups except for plant science. Plant scientists tend to prefer the term “desiccation tolerance” over “anhydrobiosis” in the literature, while animal and other researchers prefer the term “anhydrobiosis” to describe life without water. This also reflects the observation that trehalose, which has a protective role under desiccation stress, is more prevalent biologically in animals than in plants. Several of the top keywords are specific or predominantly used in the plant science literature, including the phytohormone abscisic acid, photosynthesis, and drought stress, reflecting the prevalence of plant research within the field.

**Table 1.** Top ten keywords per classification group. Terms in italics are shared across all four groups, while terms in bold are shared across the fungi, animal and microbe groups, but do not appear in the plant group.

| Kingdom | Keyword | Count |
| --- | --- | --- |
| **Plant** | *desiccation tolerance* | 557 |
|  | *desiccation* | 296 |
|  | abscisic acid | 167 |
|  | germination | 141 |
|  | cryopreservation | 121 |
|  | photosynthesis | 116 |
|  | dehydration | 104 |
|  | seed development | 98 |
|  | drought | 96 |
|  | resurrection plant | 87 |
| **Animal** | **anhydrobiosis** | 186 |
|  | *desiccation* | 174 |
|  | *desiccation tolerance* | 96 |
|  | **trehalose** | 80 |
|  | desiccation resistance | 48 |
|  | cold tolerance | 47 |
|  | tardigrada | 43 |
|  | climate change | 41 |
|  | dehydration | 39 |
|  | water balance | 35 |
| **Microbe** | *desiccation* | 70 |
|  | cyanobacteria | 55 |
|  | *desiccation tolerance* | 48 |
|  | **trehalose** | 26 |
|  | **anhydrobiosis** | 20 |
|  | biofilm | 14 |
|  | nostoc flagelliforme | 14 |
|  | photosynthesis | 14 |
|  | survival | 13 |
|  | compatible solutes | 11 |
| **Fungi** | *desiccation tolerance* | 28 |
|  | saccharomyces cerevisiae | 25 |
|  | lichens | 19 |
|  | **anhydrobiosis** | 19 |
|  | *desiccation* | 17 |
|  | **trehalose** | 17 |
|  | yeast | 12 |
|  | chlorophyll fluorescence | 12 |
|  | lichen | 11 |
|  | dehydration | 8 |

We also see these disciplinary differences present when we visualize the keywords as a network, where node size is the keyword’s frequency, and the edge weights represent how often two keywords co-occur in the same papers (Figure 2d). ‘Desiccation’, ‘desiccation tolerance’, ‘anhydrobiosis’, trehalose’, and ‘germination’ are the most interconnected keywords (Figure 2d). The preference of plant biologists for avoiding the term “anhydrobiosis” is visible in the network, as trehalose and anhydrobiosis are rarely connected to the numerous plant specific keywords related to seed development, germination, photosynthesis, resurrection plant, and abscisic acid. Similarly, “cryptobiosis”, a term specific to non-plant desiccation research, is heavily connected to “anhydrobiosis”. Interestingly, “Phenotypic plasticity” is among the most poorly connected nodes in the network, reflecting the broad view that desiccation tolerance is a binary trait, and that variability is poorly studied (Marks et al., 2021).

### Topic modeling reveals disciplinary differences in research trends

While keywords provide a general overview of research trends, they fail to offer detailed insights into specific topic areas. To address this, we employ unsupervised topic modeling, which enhances the nuance of our representations and organizes documents into distinct topic clusters. Topic modeling follows three major steps: (1) creating an embedding representation of each document, which captures the semantics, or meaning, of each document; (2) clustering document embeddings to form topic clusters; and (3) generating topic representations based on words present in each cluster. Traditional representation models generate a short list of the most important words in each topic cluster that are similar to key words, so we instead used ChatGPT-based representations with BERTopic to generate more descriptive, sentence-level topic representations.

BERTopic identified 46 topics within the desiccation tolerance literature, and the topics span from molecules to ecosystems (Figure 3a, Table S1). Topics are unevenly distributed across study systems, and few topics contain literature from animals, plants, fungi, and microbes (Figure 3b). Most topic clusters are dominated by plant literature, but 7 are dominated by animals, 3 by microbes, and 2 have predominantly fungi and microbe literature (Figure 3b). Topics with the most papers are related to environmental stress in Drosophila, understanding the molecular mechanisms of desiccation tolerance in plants, and the viability and cellular structures during desiccation in plant seeds and embryos (Table S1). Topic representations are loosely generated based on the terms that are most important for a group of documents. Study system drives the differentiation of topic clusters, rather than common methods or molecular mechanisms that are shared across groups. Interestingly, topic model 12 is the only topic with similar representation in the plant, fungi, animal, and microbe literature, and it is related to dehydration and rehydration studies in yeast. For these papers, yeast are probably used as a heterologous system for functionally validating findings from animals and plants.

**Figure 3.**
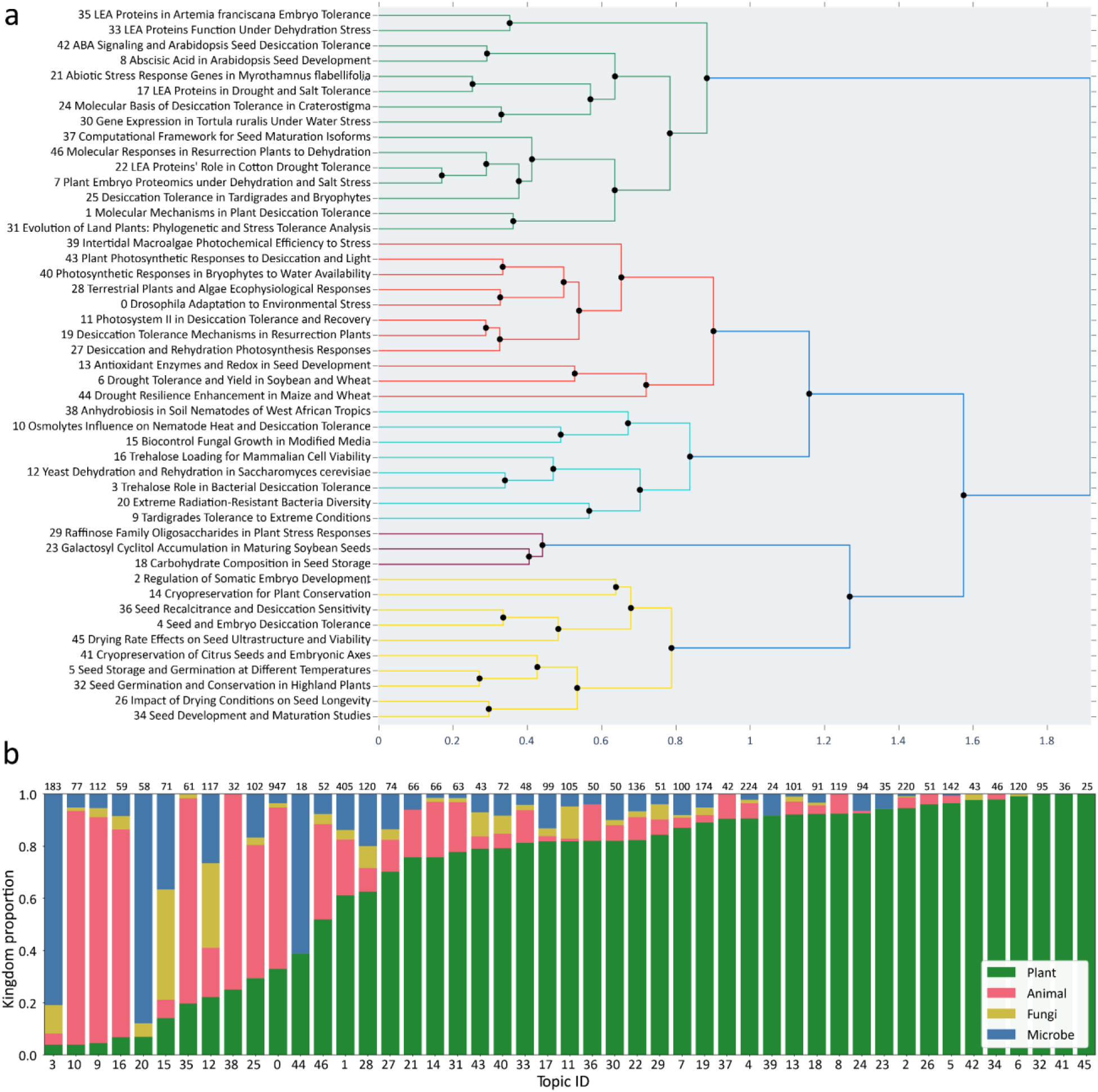
Topic models broadly separate the desiccation tolerance literature by study organism. (a) The 46 topic models are grouped into higher-order hierarchical clusters based on similarity of the ten most important terms in each cluster. Topic models are labeled by number and simplified descriptions are provided for each topic. The full ChatGPT representations for each topic ID can be found in Table S1. (b) The proportion of papers in each of the four classification groups is plotted for the 46 topic models. The number of papers belonging to each topic model is indicated above the bars.

We used hierarchical clustering to group the individual topic models into broader meta-topic representations. If we look at the meta-topic representations and the proportions of papers from each kingdom in the meta-topics (Figure 3a), we see that the hierarchical clustering groups topics that tend to study the same organism. The topic representations from hierarchical topic modeling are token-based, meaning that they are the top 5 most important words for the hierarchically-clustered group of documents. While the token-based representations are more difficult to interpret semantically, we can make more direct interpretations of the topic representations than we can for the ChatGPT-based representations.

### Citation network reveals disciplinary silos of desiccation tolerance research

Our topic modeling results heavily imply the existence of disciplinary siloing, where papers cluster together based on their study system. However, topic modeling only tells us about the content of abstracts, which typically focus on narrow findings from often siloed study systems. Given that desiccation tolerance is governed by similar molecular and biochemical mechanisms across life, insights from one system can often be applied to phylogenetically diverse species. This raises an important question: are researchers adequately citing studies from a broad range of systems?

We can visualize the connectivity of the research landscape of desiccation tolerance by building a citation network among all papers in the dataset, and coloring them by study system (Figure 4a). Nodes in the citation network represent individual papers and connections (edges) reflect co-citation, with the largest nodes having the most citations. We observed strong disciplinary siloing, where plant, animal, microbe, and fungi papers form four sparsely connected and distinct modules within the citation network (Figure 4a). The most highly cited papers tend to be influential review papers predominantly focused on desiccation tolerance in plants with one review on desiccation tolerance in prokaryotes and a mechanistic paper on LEA protein functions in animals (Table 2).

**Figure 4.**
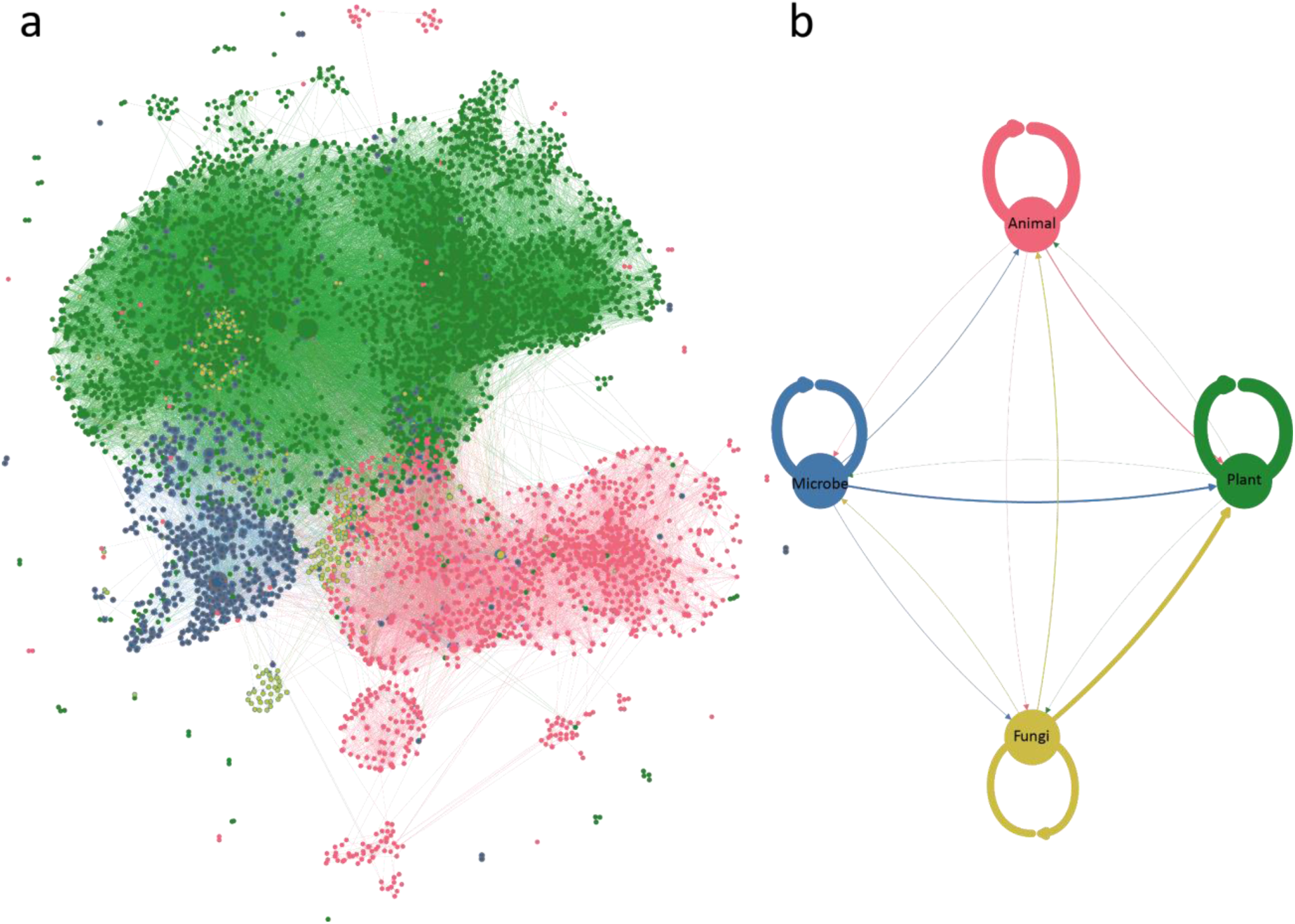
Disciplinary siloing in desiccation tolerance research. (a) Classified citation network. Each node represents a unique paper in the corpus and edges are directed citations. Nodes are colored by study system and sized by how many times they are cited, and edges are colored by the node making the citation (i.e., source node). (b) Dyadic citation frequencies. Nodes represent study groups and edges indicate one group citing another. Edges are weighted by the proportion of times one discipline cites another out of all its total citations; the thickness of all edges added together are the same for each study system group. Values for edge thickness can be found in Table S2.

**Table 2.** Ten most highly cited papers over all time in desiccation tolerance.

| Title | Authors | Year | Kingdom |
| --- | --- | --- | --- |
| Mechanisms of plant desiccation tolerance | FA Hoekstra, EA Golovina, J Buitink | 2001 | Plant |
| The molecular basis of dehydration tolerance in plants | J Ingram, D Bartels | 1996 | Plant |
| Desiccation tolerance of prokaryotes | M Potts | 1994 | Microbe |
| The evolution of vegetative desiccation tolerance in land plants | MJ Oliver, Z Tuba, BD Mishler | 2000 | Plant |
| A review of recalcitrant seed physiology in relation to desiccation-tolerance mechanisms | NW Pammenter, P Berjak, P | 1999 | Plant |
| Maturation proteins and sugars in desiccation tolerance of developing soybean seeds | SA Blackman, RL Obendorf, AC Leopold | 1992 | Plant |
| LEA proteins prevent protein aggregation due to water stress | K Goyal, LJ Walton, A Tunnacliffe | 2005 | Animal |
| Desiccation-tolerance in bryophytes: a review | MCF Proctor, MJ Oliver, AJ Wood, P Alpert, LR Stark, NL Cleavitt, BD Mishler | 2007 | Plant |
| Desiccation tolerance in bryophytes: A reflection of the primitive strategy for plant survival in dehydrating habitats? | MJ Oliver, J Velten, BD Mishler | 2005 | Plant |

We quantified disciplinary siloing by calculating dyadic citation frequencies, or the proportion of citations directed to each of the other study organism groups. Dyadic citation frequencies confirm the qualitative observation of siloing, showing that disciplines predominantly cite within their own taxonomic group (Figure 4b, Table S2). However, there are marked differences in the citation frequencies of each discipline. Plant science researchers are the most citationally isolated, citing within their own discipline 94.3% of the time. Fungal researchers are the most citationally interdisciplinary, citing their own discipline only 52.7%. However, much of the fungi literature is focused on desiccation tolerance in lichen, which contain photobionts like cyanobacteria and algae, so it is not surprising that 36.6% of the citations in fungi papers are related to plants. Notably, all other disciplines cite the plant literature as their second largest category, and animal, microbe, and fungi research communities do not cite one another nearly as often as they cite the plant literature.

While it is not possible to define a true expected baseline for an ideal world scenario for citation insularity, we can examine trends over time to see if scientists are generally becoming more or less citationally insular (Figure 5). In general, disciplines are either similar or only slightly less insular than they ever have been, with the exception of the fungal research community, which has been increasing insular over time. Increasing insularity is potentially explained by the increased rate of publishing over time. As the field of desiccation tolerance grows, it is increasingly difficult to keep up with the magnitude of information on increasingly complex discoveries, and researchers are more likely to cite within their own discipline. However, we also see that the Plant community has experienced a slight decline in insularity over the same time period, with a much larger increase in number of papers, so the potential causality of paper number driving citation insularity is not explanatory in all cases.

**Figure 5.**
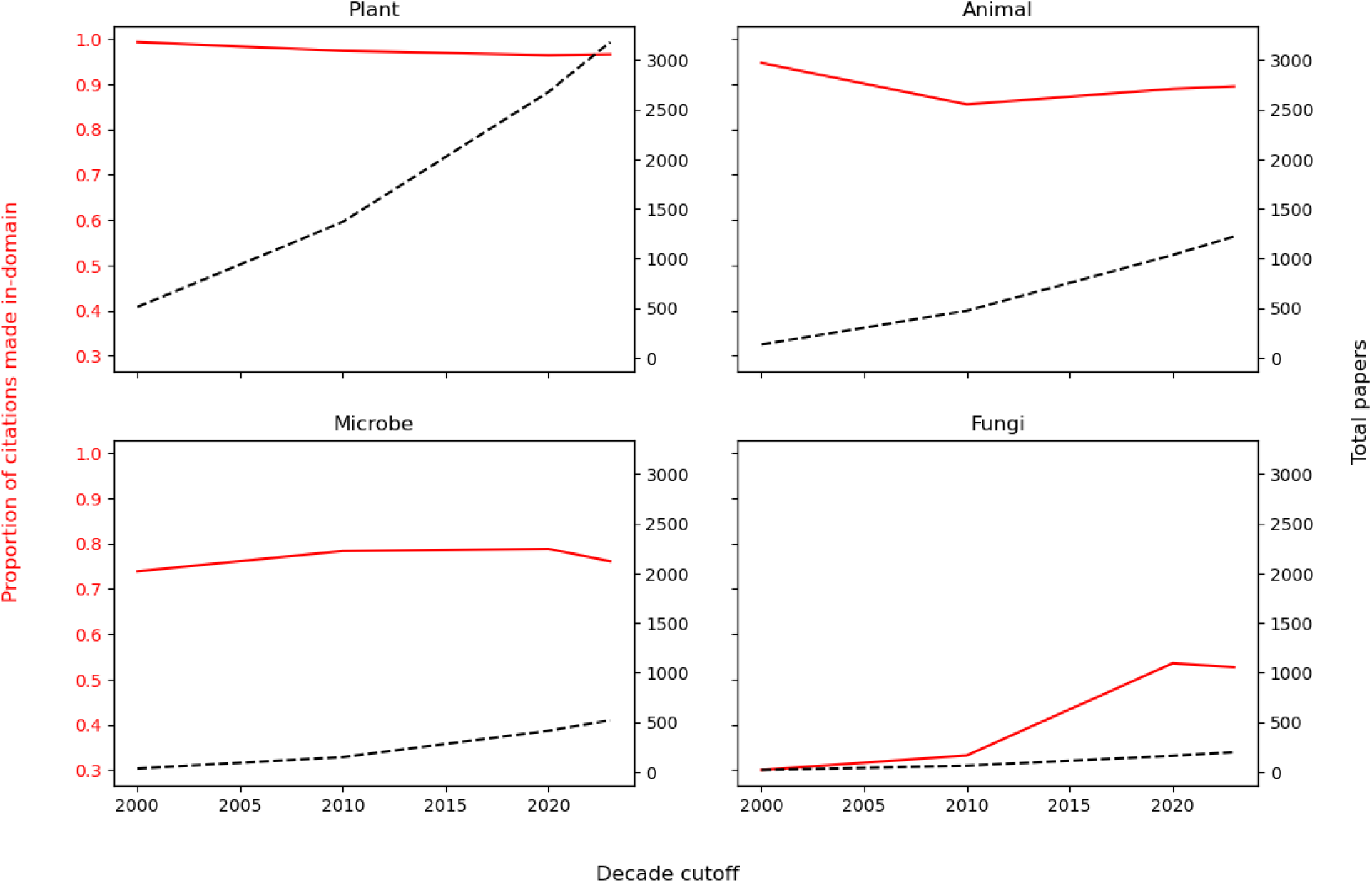
Changes in citation insularity over time. Citation insularity, or the proportion of citations made within the same taxonomic group is plotted by year along with the total number of papers for plant, animal, microbes, and fungi. Citation insularity is indicated with a solid red line (left y-axis), and total number of papers is indicated with a dotted black line (right axis).

### Citations across geography show better integration than across taxonomic groups

Given the significant siloing by discipline, we explored the relationship between geography and citation patterns. In the plant sciences, researchers from the Global South are cited half as much as those from the Global North, even when controlling for factors such as journal impact factor (Marks et al., 2023). However, we hypothesized that the field of desiccation tolerance might exhibit more balanced geographical citations due to the ease of exhaustively citing the relevant literature in a smaller field. Analyzing the citation network colored by the continent of the corresponding author reveals improved geographical integration (Figure 6a). Our graph layout algorithm does not consider the study system or continent of the nodes when arranging the graph, so the more well-mixed citation network seen in Figure 6a indicates that the groups in the graph are in fact better integrated.

**Figure 6.**
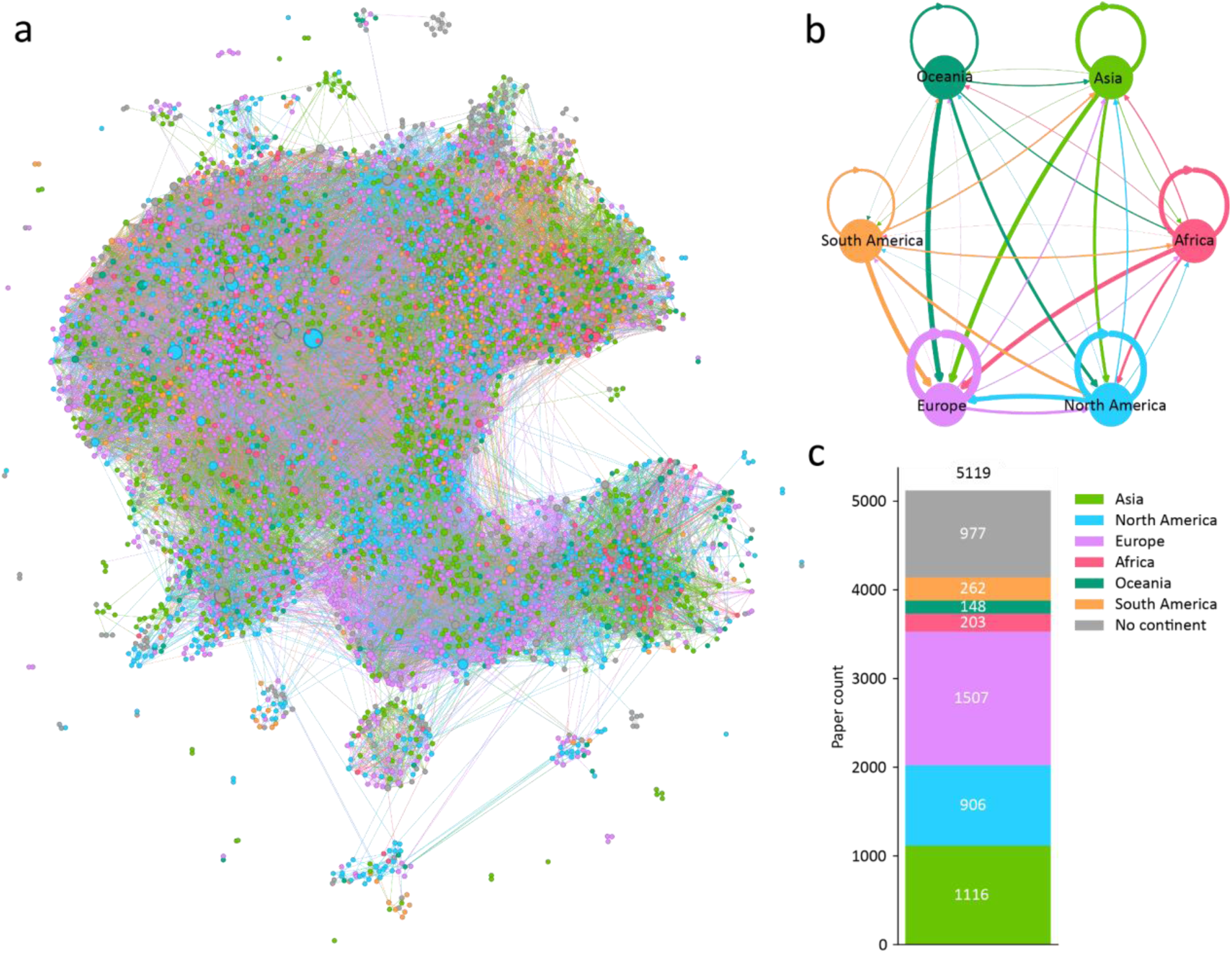
Citation insularity by paper origin. (a) The citation network from Figure 4a is recolored by the continent of the corresponding authors. (b) Dyadic citation frequencies by continent. Values for the edges can be found in Table S3. (c) Number of papers from each continent. About 19% of papers did not have geographic information recorded in the WoS database in a way that allowed us to reliably identify the continent of affiliation for the paper. Papers without a geographic category were excluded from the analysis in (b) to avoid confounding the results.

In addition to looking at the overall patterns in the citation network, we quantified continent-based siloing with the same dyadic citation analysis, both static and over time by continent (Figure 6b). While the siloing is much less severe than it was for study systems, it certainly does exist. Specifically, North America and Europe have the highest rate of self-citation, followed by Asia and Africa. However, North America, Asia, and Africa cite Europe about as often as they cite themselves, while Oceania and South America cite Europe more often than they cite their own papers. North America follows Europe as the most cited by other continents. We can ask the same question that we asked of the study system analysis: is citation insularity by continent changing over time? Figure 7 shows that all continents have gotten more insular as they have published more papers in the field of desiccation tolerance, similar to what we observed for the Fungi papers (Figure 5).

**Figure 7.**
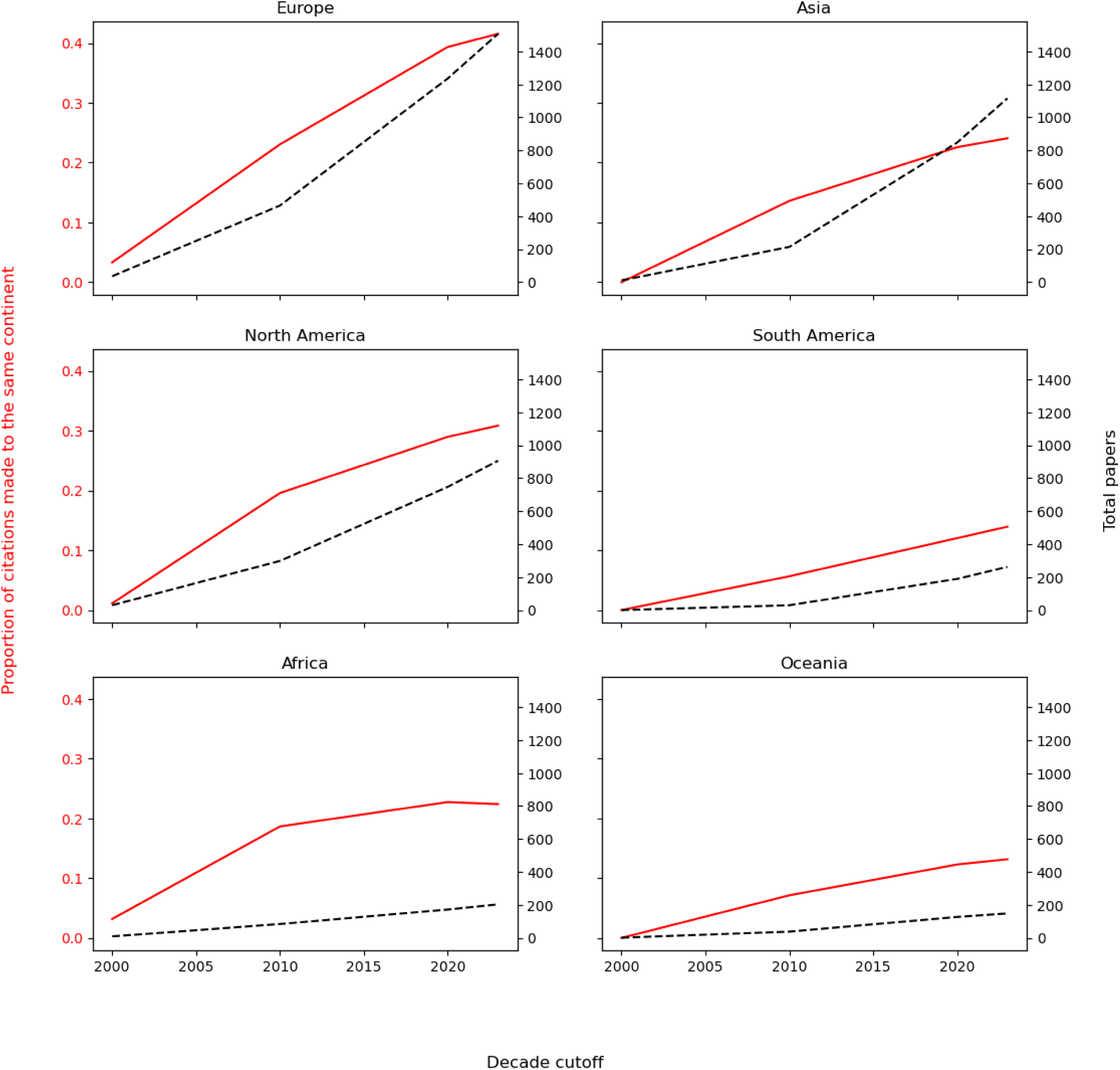
Citation insularity by continent over time. Citation insularity and paper counts are shown for Europe, Asia, North America, South America, Africa, and Oceania.

### Publication representation in biodiversity hotspots of resurrection plants

Desiccation tolerant organisms are found in virtually every biome across all seven continents, but the richest diversity of species are found in regions with seasonal drying. Resurrection plants have the highest abundance and richest diversity within the Global South, and we tested if these regions are well-represented in the literature. We obtained data on recorded observations of angiosperm resurrection plants (Table S4) and examined the number of unique species in each country (Figure 8a). We normalized the raw number of unique species in each country by the surface area in each country to control for the effect of the species-area law (Rosenzweig, 1995), which states that the more area is explored, the higher species richness will be found.

**Figure 8.**
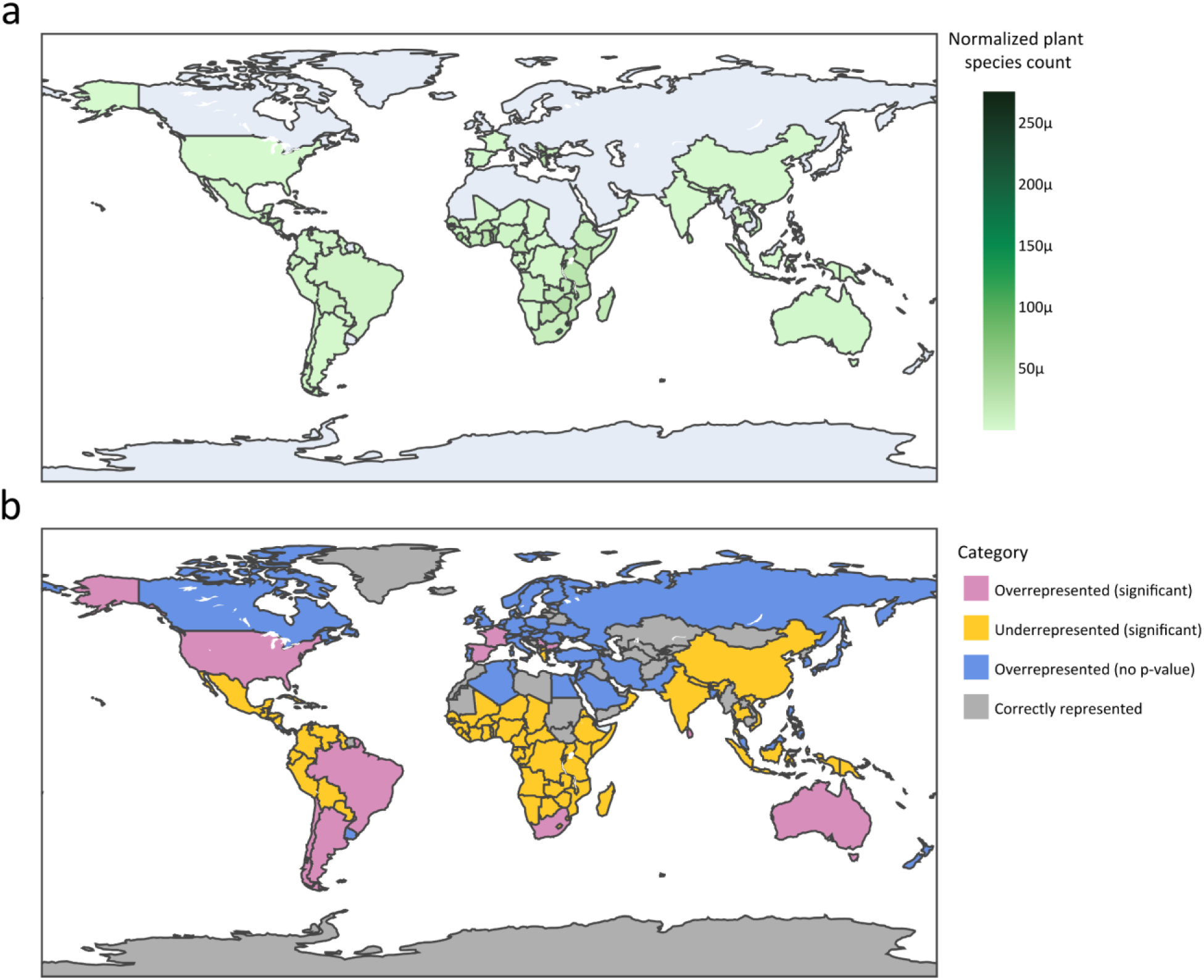
Resurrection plant distribution and literature representation. (a) Geographic distribution of unique resurrection plant species. Countries are shaded by the number of unique angiosperm resurrection plants that have been recorded within the country, normalized by the surface area of the country. Total observations for each species in each country can be found in Table S4. (b) Comparative representation of countries in the literature based on resurrection plant diversity. Chi-squared tests assessed whether the number of papers from each country (normalized by population) matched the expected number based on recorded resurrection plant species, and p-values determined significance. Countries in red were significantly overrepresented, those in purple were overrepresented because they had 0 observations of resurrection plants, but had papers in plant desiccation tolerance, countries in yellow were significantly underrepresented, and countries in grey were accurately represented, either because they did not have a significant p-value in the chi-squared, or because they had both no papers and no plant observations.

To assess the representation of countries with many resurrection plants in the literature, we first established assumptions about the expected relationship between geography and publications. A straightforward assumption is that countries with more resurrection plants should have more publications. However, population also influences publication output, as countries with more scientists generally produce more papers. Therefore we compared the number of publications, normalized by population, to the number of unique resurrection plant species recorded in each country using raw species counts rather than scaling by area, as shown in Figure 8a. Only a few countries have a representation comparable to what we would expect (Figure 8b). Contrary to our initial expectation that North American and European countries would dominate, South Africa and South American countries are also overrepresented compared to the number of resurrection plant species they contain. South Africa is a global leader in desiccation tolerance research, but much of the rest of Africa is underrepresented, despite sharing many species with South Africa, reflecting fewer publications from these regions.

### Co-citation and co-author networks reflect known interpersonal relationships

To examine relationships among authors in the desiccation tolerance literature, we analyzed their co-citation and co-author networks. Both are undirected networks; co-citation networks connect individuals based on mutual citations, while co-author networks link individuals by their joint publications. We visualize these networks for the top 3% of authors by publication number in Figures 9 and 10, respectively.

**Figure 9.**
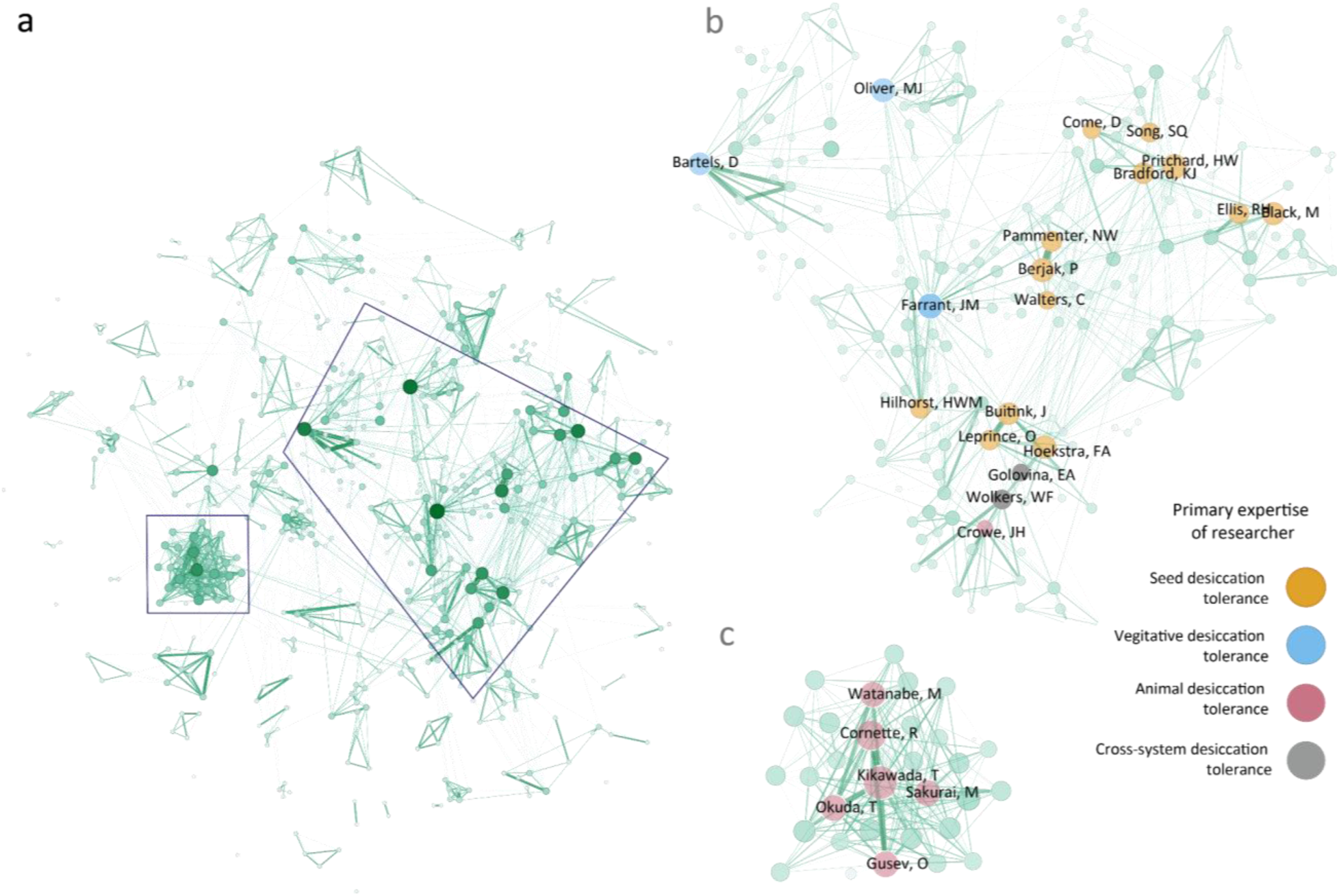
Co-authorship network of desiccation tolerance researchers. (a) Co-author network for the top 3% most productive authors. Nodes are authors and edges represent two authors collaborating together on a paper. Nodes are sized and colored by how often the author collaborates, and edges are sized and colored by how often a specific collaboration has occurred. (b) A zoomed-in subset of the co-author network demonstrating the relationships between several prominent plant desiccation tolerance researchers. (a) A zoomed-in subset of the co-author network of a highly connected cluster of microanimal researchers. This cluster exhibits a substantially higher connectivity than the rest of the co-author network, indicating that these individuals frequently co-author papers together.

**Figure 10.**
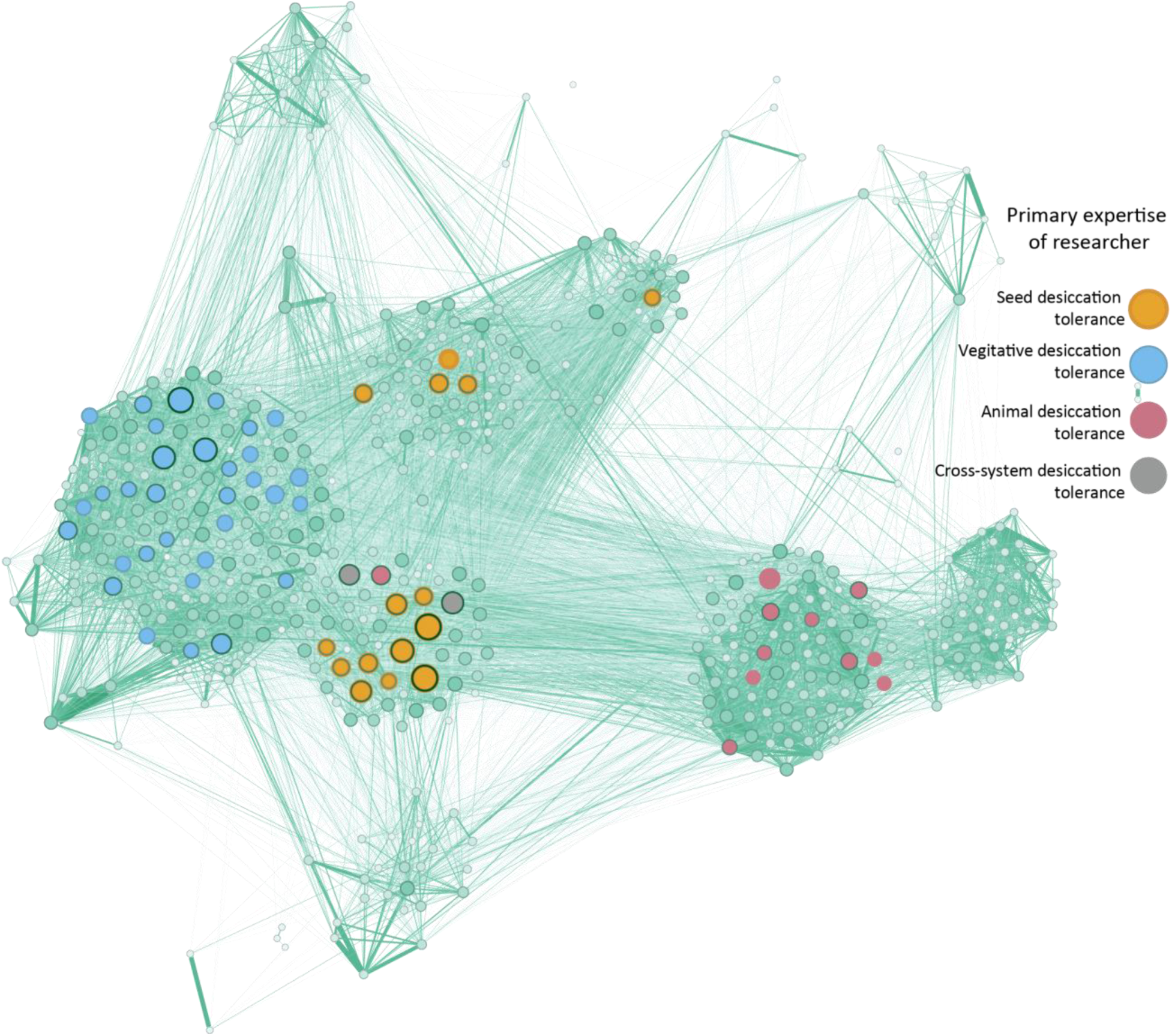
Co-citation network of desiccation tolerance experts. Co-citation network for the top 3% most productive authors is shown. Select nodes representing the most prolific authors are colored by primary study organisms.

The co-author network reveals two main hubs of highly interconnected researchers (Figure 9). The largest hub includes researchers focused on plant desiccation tolerance, with clusters on seed biology and resurrection plants. This hub also features John Crowe, a pioneer in desiccation tolerance, and cross-disciplinary researchers studying animal, fungal, microbial, and plant systems. South African plant physiologist Jill Farrant is a key central hub, connecting seed and resurrection plant biologists. Jill Farrant has a remarkably well-distributed co-author network (Figure 9b), which indicates that she collaborates with a wide group of researchers, rather than repeatedly with the same few individuals. The second hub consists of researchers on anhydrobiosis primarily in micro animals, showing high internal connectivity but sparse connections to other network researchers (Figure 9c). This group displays a remarkable amount of co-author connectivity, where almost every member of the group is connected to most others, which is a marked contrast to the more predominant patterns of either a few consistent collaborators or a hub individual that collaborates with many others separately.

To get a broader perspective on academic interactions, we examined a co-citation network using the same sets of authors (Figure 10). We see similar patterns to those observed in the co-authorship network, as individuals who collaborate are likely to work in the same discipline and study system, and cite each other heavily. Notably, individuals without direct co-authorship connections appear closely linked in the co-citation network. This is particularly evident among researchers studying resurrection plants, who frequently co-cite each other’s work despite not having published together. Additionally, since there are so many more co-citation links than co-author links, we see several distinct clusters emerge in the co-citation network as opposed to the co-author network. The resurrection plant community forms a very large, and heavily insular cluster, and the seed biology community forms three distinct but interconnected clusters. Similar to the co-authorship network, researchers working on animals form a highly connected cluster that is distinct from the plant communities.

### Conference invitee recommendation algorithm suggests candidates with related but novel expertise

To proactively utilize our bibliometric findings, we developed a rule-based algorithm to recommend new invitees to the specialized desiccation tolerance conference, DesWorks. While a supervised machine learning approach, such as XGBoost (Chen and Guestrin, 2016), could emulate past invitation processes, we opted for a heuristic approach to leverage bibliometric data differently. Our algorithm calculates a similarity score (0 to 1) based on hierarchical clustering distances in co-citation and co-author networks, topic modeling results, and geographic affiliations. This score balances novelty with relevance, penalizing individuals whose research overlaps significantly with previous attendees. We focused on the top 10 candidates, aligning with a conference capacity of 80 participants.

Our results show that even the simplest implementation of the algorithm can reveal valuable potential conference attendees. One way to evaluate these results is to look at how much of the network structure of the co-author and co-citation networks are represented by previous conference attendees (Figure 11a and c), versus when we include our proposed candidates (Figure 11 b and d). We see that none of the proposed attendees has ever co-authored a paper with a previous DesWorks attendee (Figure 11b), but that there is some degree of co-citation between proposed invitees and previous attendees (Figure 11d). This degree of integration is important. The proposed attendees are not so well-integrated with previous attendees that there is too little novelty for the conference to invite them, but co-citations suggest their research is of interest to other attendees. Additionally, when we add the proposed invitees, the sub-graph created by the previous and proposed attendees captures the overall network much better than the previous attendees alone (Figure 11b and d).

**Figure 11.**
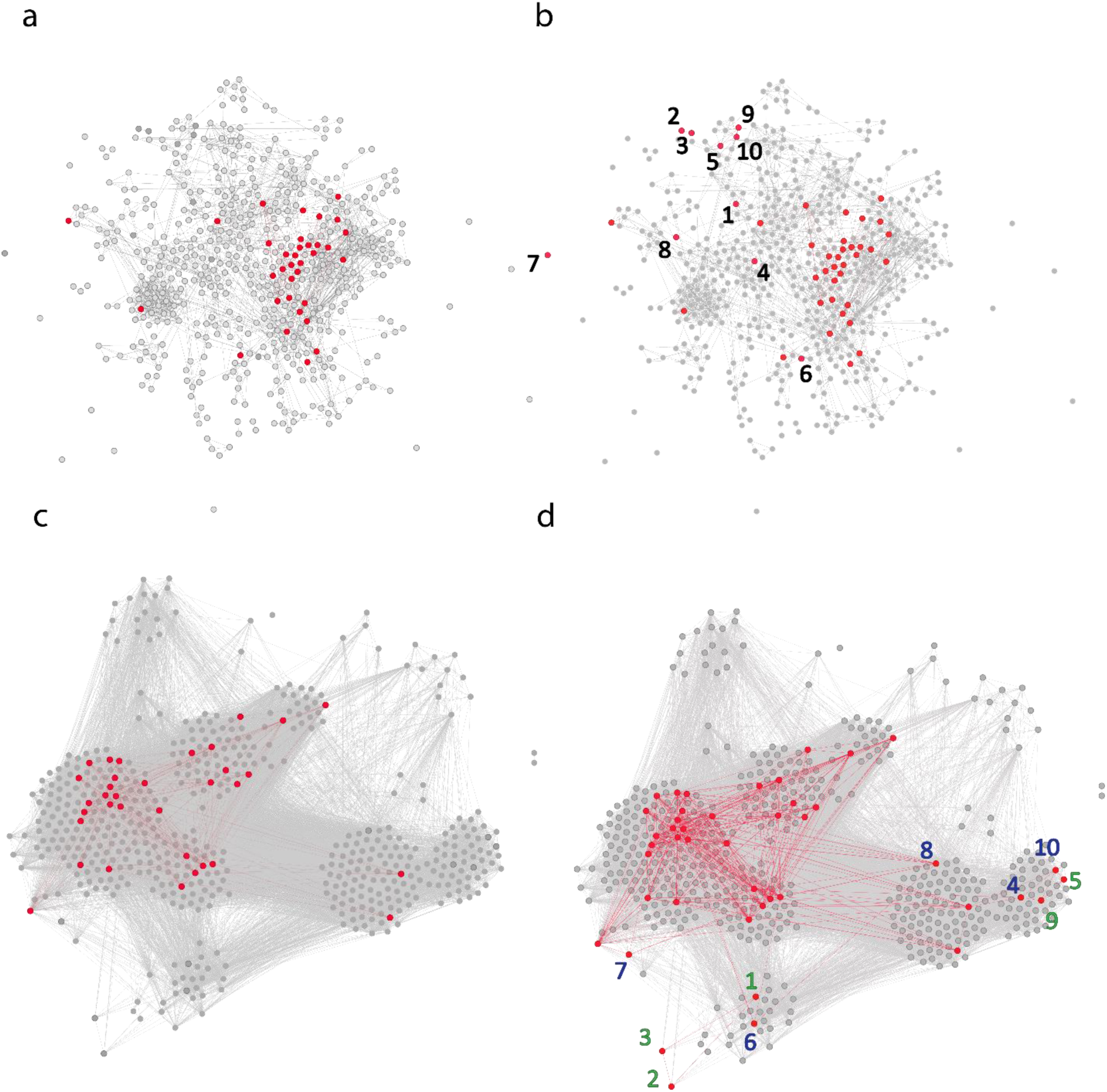
Effect of adding proposed conference invitees on network representation. Co-author network coverage before (a) and after (b) adding proposed invitees, and co-citation network coverage before (c) and after (d) adding proposed invitees. In (a) and (c), former conference attendees are colored red, with the connections between them also colored in red. In (b) and (d), proposed new attendees and their connections to one another and previous attendees are colored in red; arrows indicate proposed new attendee nodes. Green arrows indicate that the node has no direct connections to previous conference attendees, while blue arrows indicate that the node does have direct connections to previous attendees. Numbers on the arrows correspond to the candidate numbers in Table 3.

Additionally, the research specialties of the candidates (Table 3) span a diverse range of topics and include various life forms, with only two focusing exclusively on plants. This lack of plant-focused specialties indicates the algorithm’s success, as plant science is already the most represented group at DesWorks and in the broader literature. While there are many possible additions and optimizations that could be performed on our heuristic algorithm, its performance with the simplest approach is already quite impressive. We do find that authors that frequently co-author or co-cite together will receive identical scores. For example, authors 1 and 2 and authors 6 and 7 are co-authors on nearly all papers for each pair included in our dataset. However, these cases are clearly evident when looking at the interpreted results, and more candidates can be requested to fill in for essentially “duplicated” candidate entries.

**Table 3.** Candidate recommendations to attend the DesWorks conference and their research specialties.

| Candidate | Composite Score | Research Theme |
| --- | --- | --- |
| 1 | 1 | Structure and function in legume symbionts and its relation to desiccation tolerance |
| 2 | 1 | Structure and function in legume symbionts and its relation to desiccation tolerance |
| 3 | 0.9987 | Biofilm-related abiotic stress tolerance in *Listeria* |
| 4 | 0.994 | LEA proteins in plant desiccation tolerance |
| 5 | 0.9881 | Biochemistry of intrinsically disordered proteins in predominantly animal systems |
| 6 | 0.9842 | Anhydrobiosis in midges and other animals |
| 7 | 0.9842 | Anhydrobiosis in midges and other animals |
| 8 | 0.9618 | Genetics of desiccation tolerance in plants |
| 9 | 0.9375 | Drought and desiccation stress in insects |
| 10 | 0.9375 | Abiotic stress tolerance in algae |

## Discussion

In this study, we analyzed the topics and citation trends in desiccation tolerance research spanning from the 1950s to the present. DT is observed across all major lineages of life, yet much of the seminal research has been conducted in plant systems, reflecting the crucial role of seed biology in agriculture and human health. This focus contrasts with many biological fields where important discoveries were made in animal or microbial models and later applied to other systems. Our bibliometric analyses confirm that the bulk of DT research centers on plant science, though there has been a noticeable increase in contributions from animal, fungal, and microbial research systems over time. While many papers focused on model systems, we also observed a rich diversity of different resurrection plants, anhydrobiotic micro animals, and terrestrial algae and cyanobacteria within the literature, underscoring the breadth of organisms that can offer insights into desiccation tolerance. This variety not only enriches the field but also paves the way for broader discoveries across different biological systems and scales (Marks et al., 2021).

We observed strong disciplinary siloing in desiccation tolerance (DT) research, with authors predominantly citing work from similar study systems. Most topic models we identified were dominated by specific systems, with very few topics having representation from plant, fungi, animal, and microbe literature. Interestingly, we noted that larger disciplines, especially in plant sciences, tend to cite within more frequently, exhibiting increased citation insularity as they grow. This observation contradicts (Fitzpatrick et al., 2018), who proposed that smaller disciplines are more likely to cite within. Given the increasing siloing in DT research, it is essential to define our expectations and goals for interdisciplinary integration to foster scientific advancements. Future research should aim to establish a baseline for ideal citation network integration, where all relevant papers are properly cited and individual findings are placed within the context of the broader field. However, establishing such a baseline is challenging because researchers naturally gravitate towards literature within their own discipline, and the corpus of knowledge is continually expanding in scope and complexity.

When evaluating the representation of each country in the desiccation tolerance literature based on the number of resurrection plants, it’s important to acknowledge that our understanding of these plants and their geographic distribution is incomplete. Factors like research funding and international conflict unevenly distribute resources for plant surveying and collection, and there are likely many undiscovered or under-surveyed resurrection plants. Notably, countries in the Global South such as South Africa, Brazil, Argentina, and Chile appear overrepresented in the literature. This suggests that a few active researchers can significantly impact the visibility of their regions in niche scientific fields. In contrast, such overrepresentation is not observed in the broader plant sciences, which involve a large, complex, and systematically biased global researcher community (Marks et al., 2023).

Scientific conferences can enable the establishment of diverse, cross-disciplinary collaborations that can help overcome disciplinary silos and lead to new research directions. We developed a simple algorithm to recommend new attendees for specialized academic conferences based on citation metrics and author information, demonstrating its utility at the DesWorks conference. This algorithm successfully expanded the co-author and co-citation network coverage of the conference by recommending novel-yet-relevant attendees. Traditionally, the process of selecting conference attendees is done by a select group of organizing scientists, which can introduce bias in research topics and representation along various axes of diversity (Schroeder et al., 2013; Handley et al., 2015). Our algorithm offers a heuristic approach rather than a supervised machine learning model, which avoids replicating existing biases in invitation processes. However, our algorithm has limitations due to its simplicity and the constraints of available data, which might entrench existing sociological issues. It considers only four pieces of information, with two (co-citation and co-authorship) tied to publication frequency, which can be influenced by the economic status of a researcher’s country of origin. Our method also does not account for social factors like race and gender, due to the challenges and ethical concerns of making assumptions based on author names. Despite these constraints, the algorithm has significant potential for practical application. Our list of recommended attendees include new research topics and study organisms not previously represented at the DesWorks conference. We have made the code available on GitHub for further development and improvement by any research community. Our work illustrates how citation data can enhance the integration of scientists through conference participation, and we encourage more organizers to adopt such tools.

## Methods

### Data collection and preprocessing

We performed a Web of Science Core Collection search for the query “desiccation tolerance OR anhydrobiosis” on December 13, 2023, and exported 7,312 search results in the Fast 5000 format. Paper metadata, including abstracts, were collected using Michigan State University’s Web of Science Core Collection XML dataset, and some data are derived from Clarivate™ (Web of Science™). (© Clarivate 2023. All rights reserved.). The following fields were extracted for all papers: unique ID (UID), title, abstract, publication year, authors, author affiliations, references, static keywords (derived from the journal), dynamic keywords (derived from a yearly clustering analysis by Clarivate), and author-defined keywords. Because the XML dataset is a static snapshot delivered twice a year, 312 of the initial 7,312 paper UID’s did not appear in our version of the XML dataset, and were dropped. An additional 97 papers were dropped because they did not have the language tag “English” in the Web of Science database, leaving a total of 6,903 papers in the resulting dataset.

A manual examination of sample abstracts revealed the inclusion of non-biology papers from fields such as soil science, materials science, and engineering. To refine the dataset, we evaluated the potential of the dynamic and static keywords to serve as thresholds for paper filtering. There are two components of using keywords for dataset filtering: relevant/irrelevant labels for individual keywords, and the proportion of keywords for each paper that fall into the relevant/irrelevant categories. Two annotators labeled all keywords in each group as relevant or irrelevant, with agreement measured by Cohen’s kappa. The task was challenging due to the requirement for annotators to label keywords without context, resulting in low agreement (Cohen’s kappa of 0.10 for dynamic and 0.38 for static keywords). We tested both “inclusive” and “exclusive” annotations for each category.

For filtering, we considered three strategies based on the proportion of relevant keywords per paper: (1) all keywords must be relevant, (2) more than half must be relevant, or (3) at least one must be relevant. To evaluate the combinations of keyword annotations and proportional thresholding, we manually annotated 3% of the abstracts in the 6,903 paper set (∼200 abstracts) as relevant or irrelevant, with three individuals completing annotations on the same documents. Similarly to keyword annotation, each annotator applied a different level of stringency in their annotations, so agreement was low (0.30, 0.17, and 0.13 between each pair of the three annotators). Document selection at the early stages of any pipeline that analyzes the literature biases the downstream analyses performed on the resulting dataset. Therefore, we decided to test all thresholding approaches against the most inclusive version of the annotated test set. There were two relatively equally well performing strategies on our test set: both static and dynamic keywords annotated by the more inclusive annotator, using the least stringent filtering strategy where only one keyword needed to be relevant (F1 scores of 0.90 and 0.93, respectively). Using static keywords resulted in ∼250 fewer papers in the filtered dataset than when using dynamic keywords, so we chose the static keyword-least stringent approach to balance inclusivity with exclusion. The final dataset of DT literature contained 5,963 papers.

### Citation network construction and paper classification

We built a directed citation network, where nodes are papers and directed edges are citations, and classified each node according to the kingdom of its study organism. Following (Fitzpatrick et al., 2018), we only consider citations to papers that are also included within the main search result dataset of 5,963 papers. This approach was chosen due to the computational complexity of obtaining and classifying abstracts and other metadata for all references. Additionally, many references lack complete information, such as UIDs, making it challenging to determine their validity automatically. We also excluded papers without an abstract recorded in the XML dataset, resulting in the removal of 337 papers and leaving a total of 5,626 papers in the citation network.

We defined four study system groups based on the taxonomic classifications in NCBI Taxonomy: Plant (*Viridiplantae*), Animal (*Metazoa*), Microbe (*Bacteria* and *Archaea*), and Fungi (*Fungi*). To classify papers into groups, we first used the TaxoNERD version 1.5.2 classifier (Le Guillarme and Thuiller, 2022) to identify the scientific and common name mentions of each species, and link them back to the NCBI Taxonomy database, allowing us to classify each species into a taxonomic group. For review or biochemistry papers where no species were identified, we used fuzzy string matching against a predefined mapping of generic terms to kingdoms to further classify papers. If an abstract contained a string matching at least two-thirds of the characters of a generic term, it was classified under that group.

For papers mentioning multiple species from different groups, we assigned the classification based on the most represented group in the abstract. For example, if there were five species mentioned, and three of them were in the Plant category and two in the Animal category, the paper would be classified as Plant. Classifications were manually reviewed and corrected using specific heuristics for difficult-to-classify organisms: (a) algae (Chlorophyta, Rhodophyta, and Phaeophyta, or anything referred to as red, green, or brown algae) were considered plants; (b) lichens were considered fungi; and (c) diatoms, non-photosynthetic protists, and viruses were considered microbes. Papers mentioning multiple symbiotic organisms were classified based on the focal organism; for example, a paper studying the microalgal symbiont of a fungi was classified as Plant as opposed to a Fungi paper, while a paper studying the effect of a biocontrol bacteria on a plant was classified as a Microbe paper. If a paper studied separate organisms from multiple kingdoms in relatively equal numbers (e.g. mosses and lichens, or cyanobacteria and eukaryotic microalgae), we use the grouping that best reflects how the organisms were studied. For example, a paper that studies cyanobacteria and microalgae together in their role as microorganisms was classified as Microbe, whereas a paper that studies mosses and lichens with respect to their photosynthesis was classified as Plant.

We aimed to include as many papers as possible and maintained transparency in our classification approach during manual review. However, papers that could not fit into a category, even with some interdisciplinary ambiguity, were considered unclassifiable and removed from the dataset. Examples include pure biochemistry papers that do not list the organisms of origin for the studied proteins or ecology papers examining interactions across all kingdoms of life.

In addition to classification of papers, we used TaxoNERD to identify the number of unique species in each taxonomic group mentioned in our dataset. We removed all identified mentions above the species level (genus, family, etc) when performing the final calculations of unique mentions in each group. We also calculated the number of new and cumulative papers per year, using the publication year information obtained from the XML dataset, and split this calculation into the study system groups with the paper classifications obtained during citation network construction.

### Keyword analysis

Author-identified keywords for all papers were lowercased and separated by kingdom of their paper of origin. We visualized the number of keywords per paper with a histogram, created a seaborn density plot per kingdom to examine keyword frequency in the dataset, and used the UpSetPlot package to visualize the overlap in keywords across kingdoms. We took the top 1% of keywords across the entire dataset by frequency and built a co-occurrence network, where nodes are keywords and edges are weighted by how often two keywords appear together on the same paper. To further examine the top 1% of keywords, we plotted (a) absolute keyword frequency per year, (b) keyword frequency per year normalized by the number of new publications in that year, and (c) the absolute and normalized number of new keywords introduced per year.

### Topic modeling

To characterize the landscape of topics in the field of desiccation tolerance, we applied a topic modeling approach using BERTopic (Grootendorst, 2022). We used the SentenceTransformer model ‘allenai/scibert_scivocab_cased’ for the sentence model. For clustering, we employed UMAP with parameters set to n_neighbors=15, n_components=5, min_dist=0.0, metric=’cosine’, and random_state=42. The CountVectorizer was utilized for vectorization with settings of stop_words=“english”, ngram_range=(1, 3), and min_df=10. Finally, for representation, we used OpenAI’s “gpt-3.5-turbo” model with the chat feature enabled.

Abstracts were passed as-is (e.g., no pre-processing) to the BERTopic pipeline, as the advantage of a transformer-based topic modeling pipeline is that the model can leverage all available context, including stop words. After the initial model fitting, outliers were reduced with the “embeddings” strategy at a threshold of 0.1. Topic clusters were visualized according to the proportion of documents in each cluster that belong to each kingdom. Hierarchical topical modeling was performed with the functions “hierarchical_topics” and “visualize_hierarchy”.

### Citation network visualization and citation insularity

We performed citation network visualization and calculated citation insularity for publications for both study system groupings as well as geography. In both cases, the directed citation network was visualized using the tool Gephi (Bastian et al., 2009) with the OpenOrd layout algorithm (random seed 3171345782275371875). Nodes are colored by their study system group or the continent of their corresponding author, and are sized by their in-degree, or how often they are cited. Edges are colored the same as the node from which they originate.

Citation insularity, which measures how often authors within a discipline cite works outside their field, was examined using dyadic citation frequencies as described by Fitzpatrick et al. (Fitzpatrick et al., 2018). Prior to performing calculations, we removed all papers without a classification for study system or continent. For each directional pair of disciplines (e.g., Plant → Animal, Fungi → Microbe) or continents, we calculated the dyadic citation frequency as the proportion of citations from the first category to the second. For example, if 20% of citations in Animal papers are to Microbe papers, the dyadic citation frequency for Animal → Microbe would be 0.2. Additionally, we calculated and analyzed the dyadic citation frequencies from one discipline or continent to itself over time, which we refer to as citation insularity, and reflects how often a group cites its own work. These calculations were performed for all papers up to the end of each decade. We also calculated and visualized the total number of papers in each discipline for those dates on the same plot.

### Geographic representation of plant science papers

Data on global resurrection plant distribution was obtained via personal communication with Luiz Bondi. Coordinates for resurrection plant distributions were reverse mapped to identify their country of origin using the GeoPy package (Lopez Gonzalez-Nieto et al., 2020). After reverse mapping, we plotted the number of unique species in each country divided by the surface area (from the Natural Earth low resolution and tiny countries datasets) of that country to account for the species-area law.

The calculation of paper representation based on plant count involved the following quantities: paper count, plant count, and population of each country. We used the Natural Earth low resolution and tiny countries datasets to obtain the iso-alpha3 codes for all countries, and used the World Bank API to obtain population values from 2022 for all countries, with the exception of Taiwan which we obtained separately from a Taiwanese government website (https://www.ris.gov.tw/app/en/2121?sn=23009515). Each paper was assigned a country based on the affiliation of the corresponding author and papers that could not be classified into a country were dropped for this analysis.

For each country, we calculated an expected paper count, as in:

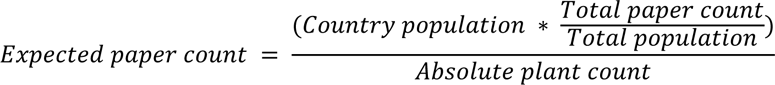

This analysis assumes that each country functions as a cultural unit, and that the number of researchers studying resurrection plants does not scale with the number of species in their local area, but rather with their cultural network. The observed paper count, against which the expected paper count is compared, is the number of publications in the dataset from that country, divided by the absolute plant count.

To determine whether the difference between the expected and observed plant counts was significant, we performed a chi-squared test loop, where the observed and expected values from each country are compared against the observed and expected values from all other countries. We performed a multiple testing correction on the resulting p-values using the scipy.stats.false_discovery_control implementation of the Benjamini-Hochberg method. Countries that had papers in the dataset but a plant count of 0 could not be tested but are considered to be overrepresented, while countries that had both 0 papers and 0 plants are considered correctly represented.

### Co-citation and co-author networks

Co-citation and co-author networks were built as part of the conference invitee recommendation algorithm code (see below). In short, the co-author network was constructed by counting the number of times that each possible pair of authors appear on the same paper. That number determines the weight of each edge in the network. For co-citation, the full citation network is processed to count the number of times that each pair of authors cites one another, which determines the weight of edges in the co-citation network. Importantly, edges in the co-citation network are undirected, and account for both directions of citation between a pair of authors. To aid clarity in visualization, only the top 3% of authors by publication productivity are kept, and only nodes with more than 20 co-authorships or more than 200 co-citations are labeled in the final visualizations.

### Conference invitee recommendation algorithm

To enhance the global reach and diversity of invitations to specialized conferences, we developed a recommendation algorithm using the Desiccation Workshop (DesWorks) conference as a case study. DesWorks, a small conference focused on desiccation tolerance, has been held every four years in South Africa since 1994. The choice of DesWorks was driven by our familiarity with the field and the historical intent of the conference organizers to diversify attendees and topics. Despite these efforts, invitations are still largely extended through personal networks of organizers and past attendees, presenting an opportunity for systematic improvement.

For this project, we obtained the 2016 and 2024 conference attendee lists by contacting the organizers. Due to limited historical data availability, our analysis began with 2016 records. We manually verified the publication names of attendees, as registration names often differed from publishing names, to consolidate their publications in our dataset. For all other authors, we assumed unique Web of Science-formatted names represented distinct individuals. All data was partially anonymized for publication.

The algorithm leverages bibliometric data to recommend potential invitees based on co-citation and co-author networks, topic modeling results, and authors’ country of affiliation. It identifies researchers who might bring novel insights to DesWorks but are only loosely connected to current attendees through citation patterns. The algorithm calculates a score for each author by assessing their proximity in the co-citation and co-author networks, novelty in topic modeling clusters, and geographic representation.

A summary of the algorithm is as follows: for each of the four elements (co-citation and co-author networks, topic model clusters, and geographic location, Algorithm 1), we calculate a score for each author that balances the similarities and differences between that author and all conference attendees (Algorithm 2). For the two networks, we calculate the mean hierarchical distance between the author and conference attendees. We want the candidate invitees to be close, but not too close, to previous attendees in the network, so we apply a calculation that privileges being closest to the 25th percentile of all author-to-attendee distances. For topic modeling, we optimize for topic novelty. The author gets a 0 if they are in the same topic model cluster as conference attendees or 1 if there are no conference attendees in their cluster. Importantly, for topic modeling and for the two network types, authors can be present in multiple clusters, so we calculate these scores for each occurrence of the author and then average them to get to the final score for each element. We aimed to expand the representation at the conference geographically, however, just because a region is represented in the conference does not mean it has proportional representation.

To account for low representation, the geographic score is calculated as 1 for a country not present in the conference, 0.5 for a country in the conference but that falls in the bottom half of countries by attendee count, and 0 if the country is in the top 50th percentile of country representation at the conference. Finally, all component scores are averaged to get each author’s score, and the top authors by score are presented as invitee candidates. The number of authors presented as candidates can be set by the user; we set ours to 10 individuals. Importantly, not all authors have geographic locations recorded in the dataset.

One major issue with the WoS XML dataset is that, while some papers have addresses recorded for the authors, they are missing an index to relate each author to their address. For papers where all addresses are in the same country, we were able to assign a location for the authors. However, 524 papers lacked a clear mapping between authors and addresses and included addresses from multiple countries, making it impossible to accurately map authors to their countries of origin. This issue was more prevalent than the 282 papers that completely lacked geographic information. Authors without a reliably identifiable geographic location are automatically assigned what is referred to as the *general probability score* in Algorithm 2. The calculation of the general probability score follows the calculation:

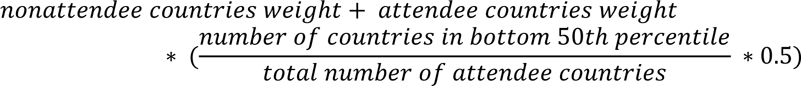

For the sake of this calculation, we assume that the actual distribution of authors in the whole dataset follows the visible distribution of authors with recorded countries, and calculate the probability that a random author belongs to either (a) a high representation country (score of 0), (b) a low representation country (score of 0.5), or (c) a country that is not represented (score of 1), and multiply this probability of membership by the score for that kind of membership. The term for belonging to a high-representation country for the conference falls out because it is multiplied by 0, so the final calculation is the sum of the probability that the individual belongs to a non-represented country plus the probability that they belong to a low-representation country.

Finally, we tested the application of an enrichment threshold. In the basic formulation of the algorithm (Algorithm 1), any cluster containing a conference attendee is considered enriched for conference attendees. However, we may want to exclude clusters that only have one or two conference attendees in them. For each type of cluster, after calculating the percentage conference attendees in each cluster (the “enrichment”), we applied a threshold by taking the median of the enrichment values, and setting the enrichment to 0 for any cluster that fell below the median. The threshold value can be set at an arbitrary percentile, but we started with the 50th percentile to measure the effect of a middle-ground threshold. We found that the 50th percentile enrichment threshold only meaningfully impacted the topic scores (Figure S1); however, that change diversified the scores for the top ten candidates, so we maintained the 50th percentile enrichment threshold in our analyses.

To characterize the output of the conference invitee recommendation algorithm, we obtained the titles, abstracts, publication years, and affiliation at the time of publication for all candidates’ publications within our dataset, and then manually characterized the themes of those publications for each candidate.

## Acknowledgements

This work was funded by NSF DBI-2213983 to RV. We thank Luiz Bondi for providing the raw data for desiccation tolerant plant observations, and Keren Cooper for providing the DesWorks attendee lists. We also thank Devin Higgins from MSU Libraries and Maureen Sico from Clarviate for their help in using the Web of Science XML dataset.

## Data availability

The code for these analyses is available on a dedicated github page: https://github.com/serenalotreck/desiccation-network.

