## Supplemental Figures/Tables for "Unifying the Research Landscape of Desiccation Tolerance to Identify Trends, Gaps, and Opportunities"

**Table S1. ChatGPT-based topic representations.**

| **Topic ID** | **Representation** |
| --- | --- |
| 0 | Adaptation mechanisms to environmental stress in Drosophila species |
| 1 | Understanding Molecular Mechanisms of Desiccation Tolerance in Plants |
| 2 | Regulation of Somatic Embryo Development through Carbohydrates, Osmoticum, and Storage Reserves |
| 3 | Role of Trehalose in Desiccation Tolerance of Bacterial Strains |
| 4 | Desiccation tolerance in seeds and embryos: impact of drying rates and water content on viability and cellular structures |
| 5 | Seed Storage Behavior and Germination in Different Temperatures |
| 6 | Influence of Soil Drying on Drought Tolerance and Yield in Different Soybean and Wheat Cultivars |
| 7 | Proteomic analysis of stress responses and metabolic regulation in plant embryos under dehydration and salt stress |
| 8 | Abscisic acid regulation in Arabidopsis seed development and response to osmotic stress |
| 9 | Tardigrades' Tolerance to Extreme Conditions and Radiation |
| 10 | Influence of Osmolytes on Heat Tolerance and Desiccation Tolerance in Entomopathogenic Nematodes |
| 11 | The Role of Photosystem II in Desiccation Tolerance and Recovery |
| 12 | Yeast Dehydration and Rehydration Studies in Saccharomyces cerevisiae |
| 13 | Changes in antioxidant enzyme activities and redox balance during seed development and desiccation |
| 14 | Cryopreservation Techniques for Ex Situ Plant Conservation |
| 15 | Biocontrol Fungal Growth and Survival in Modified Media |
| 16 | Enhancing mammalian cell viability through intracellular trehalose loading for freeze-drying and desiccation tolerance |
| 17 | Role of LEA proteins in enhancing drought and salt stress tolerance in transgenic plants |
| 18 | Soluble carbohydrate composition and storage in seeds of various plant species |
| 19 | Physiological and Metabolic Mechanisms of Desiccation Tolerance in Resurrection Plants |
| 20 | Diversity and Characteristics of Extreme Radiation-Resistant Bacteria in Various Environments |
| 21 | Plant Abiotic Stress Response Genes in Myrothamnus flabellifolia and Other Plant Species |
| 22 | Characterization and Role of Late Embryogenesis Abundant (LEA) Proteins in Drought Stress Tolerance in Upland Cotton |
| 23 | Accumulation of galactosyl cyclitols in soybean seeds during maturation |
| 24 | Molecular Basis of Desiccation Tolerance in Craterostigma Plantagineum |
| 25 | Molecular Mechanisms of Desiccation Tolerance in Tardigrades and Bryophytes |
| 26 | The Impact of Different Drying Conditions on Seed Longevity and Quality |
| 27 | Photosynthetic responses to desiccation and rehydration in various plant species |
| 28 | Ecophysiological Responses of Various Terrestrial Plants and Algae to Different Abiotic Conditions |
| 29 | Role of Raffinose Family Oligosaccharides (RFOs) in Plant Stress Responses |
| 30 | Characterization of Gene Expression and Protein Function in Tortula ruralis Under Water Deficit Stress |
| 31 | Evolution of Land Plants and Bryophytes: Phylogenetic Analysis of Early Lineages, Gene Families, and ABA-Dependent Stress Tolerance |
| 32 | Seed germination and conservation strategies for tropical highland and subtropical Andean plant species |
| 33 | Structure and Function of Late Embryogenesis Abundant (LEA) Proteins Under Dehydration Stress |
| 34 | Seed Development and Maturation Studies in Various Plant Species |
| 35 | Role of Late Embryogenesis Abundant (LEA) Proteins in Desiccation Tolerance of Artemia franciscana Embryos |
| 36 | Understanding Seed Recalcitrance and Desiccation Sensitivity |
| 37 | Integrated computational framework for predicting isoform-specific functions during plant seed maturation |
| 38 | Anhydrobiosis and Population Dynamics of Soil Nematodes in Semi-arid Tropics of West Africa |
| 39 | Photochemical efficiency responses of intertidal brown macroalgae to multiple environmental stressors |
| 40 | Impact of Water Availability on Photosynthetic Responses in Polar and Temperate Bryophytes |
| 41 | Cryopreservation techniques for citrus seeds and embryonic axes |
| 42 | Relationship between ABA signaling and desiccation tolerance in Arabidopsis seeds |
| 43 | Photosynthetic Responses to Desiccation and Light Intensity Among Various Plant Species |
| 44 | Potential of plant growth promoting bacteria for enhancing drought resilience in maize and wheat under water deficit conditions |
| 45 | Comparative study on the effect of drying rate on viability and ultrastructure of embryonic axes in recalcitrant seeds |
| 46 | Molecular Responses to Dehydration and Rehydration in Resurrection Plants |

**Table S2. Dyadic citation frequencies by study system.**

| **Source** | **Target** | **Dyadic frequency** |
| --- | --- | --- |
| Plant | Plant | 0.966 |
|  | Animal | 0.015 |
|  | Microbe | 0.01 |
|  | Fungi | 0.01 |
| Animal | Plant | 0.017 |
|  | Animal | 0.895 |
|  | Microbe | 0.017 |
|  | Fungi | 0.016 |
| Microbe | Plant | 0.151 |
|  | Animal | 0.055 |
|  | Microbe | 0.76 |
|  | Fungi | 0.034 |
| Fungi | Plant | 0.363 |
|  | Animal | 0.066 |
|  | Microbe | 0.044 |
|  | Fungi | 0.527 |

**Table S3. Dyadic citation frequencies by continent**

| **Source** | **Target** | **Dyadic frequency** |
| --- | --- | --- |
| Africa | Africa | 0.345 |
|  | Asia | 0.087 |
|  | Europe | 0.319 |
|  | North America | 0.182 |
|  | Oceania | 0.054 |
|  | South America | 0.013 |
| Asia | Africa | 0.065 |
|  | Asia | 0.333 |
|  | Europe | 0.334 |
|  | North America | 0.216 |
|  | Oceania | 0.027 |
|  | South America | 0.026 |
| Europe | Africa | 0.067 |
|  | Asia | 0.1 |
|  | Europe | 0.58 |
|  | North America | 0.212 |
|  | Oceania | 0.025 |
|  | South America | 0.016 |
| North America | Africa | 0.058 |
|  | Asia | 0.102 |
|  | Europe | 0.347 |
|  | North America | 0.447 |
|  | Oceania | 0.029 |
|  | South America | 0.017 |
| Oceania | Africa | 0.104 |
|  | Asia | 0.1 |
|  | Europe | 0.364 |
|  | North America | 0.222 |
|  | Oceania | 0.197 |
|  | South America | 0.013 |
| South America | Africa | 0.122 |
|  | Asia | 0.121 |
|  | Europe | 0.33 |
|  | North America | 0.215 |
|  | Oceania | 0.032 |
|  | South America | 0.181 |

**Table S4. Species observation counts per country.**

| **Country Code** | **Species** | **Observed Count** |
| --- | --- | --- |
| AGO | *Craterostigma plantagineum* | 2 |
|  | *Myrothamnus flabellifolius* | 1 |
|  | *Eragrostis nindensis* | 1 |
|  | *Microchloa indica* | 3 |
|  | *Microchloa kunthii* | 1 |
|  | *Micrachne patentiflora* | 1 |
|  | *Sporobolus stapfianus* | 1 |
|  | *Xerophyta squarrosa* | 4 |
|  | *Xerophyta scabrida* | 1 |
| ARG | *Blossfeldia liliputana* | 5 |
|  | *Microchloa indica* | 40 |
|  | *Microchloa kunthii* | 31 |
|  | *Tripogonella spicata* | 41 |
|  | *Barbaceniopsis humahuaquensis* | 47 |
|  | *Barbaceniopsis boliviensis* | 5 |
| AUS | *Borya constricta* | 362 |
|  | *Borya inopinata* | 6 |
|  | *Borya mirabilis* | 16 |
|  | *Borya nitida* | 18 |
|  | *Borya septentrionalis* | 74 |
|  | *Borya scirpoidea* | 46 |
|  | *Borya sphaerocephala* | 705 |
|  | *Boea hygroscopica* | 109 |
|  | *Micraira multinervia* | 26 |
|  | *Eragrostiella bifaria* | 14 |
|  | *Micraira adamsii* | 128 |
|  | *Microchloa indica* | 12 |
|  | *Micraira lazaridis* | 25 |
|  | *Micraira subulifolia* | 35 |
|  | *Micraira spinifera* | 19 |
|  | *Micraira tenuis* | 77 |
|  | *Micraira viscidula* | 24 |
|  | *Sporobolus elongatus* | 99 |
|  | *Tripogonella loliiformis* | 8 |
| BDI | *Microchloa kunthii* | 2 |
|  | *Sporobolus stapfianus* | 1 |
| BEN | *Afrotrilepis pilosa* | 3 |
|  | *Microchloa indica* | 39 |
|  | *Microchloa kunthii* | 9 |
|  | *Oropetium aristatum* | 6 |
|  | *Sporobolus festivus* | 23 |
|  | *Tripogonella minima* | 21 |
| BFA | *Afrotrilepis pilosa* | 1 |
|  | *Microchloa indica* | 22 |
|  | *Oropetium aristatum* | 13 |
|  | *Sporobolus festivus* | 12 |
|  | *Sporobolus pellucidus* | 1 |
|  | *Tripogonella minima* | 11 |
| BGR | *Haberlea rhodopensis* | 7 |
| BOL | *Pitcairnia lanuginosa* | 35 |
|  | *Blossfeldia liliputana* | 2 |
|  | *Microchloa indica* | 27 |
|  | *Microchloa kunthii* | 6 |
|  | *Tripogonella spicata* | 9 |
|  | *Vellozia variabilis* | 5 |
|  | *Vellozia andina* | 18 |
|  | *Barbaceniopsis boliviensis* | 13 |
|  | *Vellozia caruncularis* | 3 |
|  | *Vellozia tubiflora* | 12 |
| BRA | *Pitcairnia lanuginosa* | 140 |
|  | *Trilepis ciliatifolia* | 14 |
|  | *Trilepis lhotzkiana* | 133 |
|  | *Trilepis microstachya* | 3 |
|  | *Microchloa indica* | 46 |
|  | *Tripogonella spicata* | 35 |
|  | *Barbacenia blackii* | 17 |
|  | *Barbacenia fanniae* | 10 |
|  | *Barbacenia flava* | 65 |
|  | *Barbacenia fragrans* | 4 |
|  | *Barbacenia longiflora* | 24 |
|  | *Barbacenia gentianoides* | 34 |
|  | *Barbacenia graminifolia* | 7 |
|  | *Barbacenia longiscapa* | 17 |
|  | *Barbacenia macrantha* | 27 |
|  | *Barbacenia purpurea* | 17 |
|  | *Barbacenia riedeliana* | 9 |
|  | *Barbacenia seubertiana* | 7 |
|  | *Barbacenia spectabilis* | 2 |
|  | *Barbacenia tomentosa* | 8 |
|  | *Barbacenia gounelleana* | 53 |
|  | *Vellozia variabilis* | 30 |
|  | *Vellozia albiflora* | 47 |
|  | *Vellozia angustifolia* | 10 |
|  | *Vellozia candida* | 46 |
|  | *Vellozia ciliata* | 9 |
|  | *Vellozia caput-ardeae* | 14 |
|  | *Vellozia caruncularis* | 61 |
|  | *Vellozia compacta* | 35 |
|  | *Vellozia declinans* | 131 |
|  | *Vellozia epidendroides* | 258 |
|  | *Vellozia glochidea* | 26 |
|  | *Vellozia hatschbachii* | 7 |
|  | *Vellozia hirsuta* | 295 |
|  | *Vellozia nanuzae* | 27 |
|  | *Vellozia nivea* | 28 |
|  | *Vellozia plicata* | 40 |
|  | *Vellozia pulchra* | 19 |
|  | *Vellozia resinosa* | 14 |
|  | *Vellozia flavicans* | 78 |
|  | *Vellozia sellowii* | 6 |
|  | *Vellozia squalida* | 1 |
|  | *Vellozia semirii* | 15 |
|  | *Vellozia streptophylla* | 8 |
|  | *Vellozia subscabra* | 19 |
|  | *Vellozia taxifolia* | 9 |
|  | *Vellozia tubiflora* | 152 |
|  | *Vellozia variegata* | 52 |
|  | *Vellozia verruculosa* | 2 |
| BTN | *Eragrostiella nardoides* | 1 |
|  | *Microchloa kunthii* | 2 |
| BWA | *Craterostigma plantagineum* | 1 |
|  | *Eragrostis nindensis* | 4 |
|  | *Microchloa caffra* | 1 |
|  | *Microchloa indica* | 2 |
|  | *Microchloa kunthii* | 1 |
|  | *Micrachne patentiflora* | 2 |
|  | *Oropetium capense* | 10 |
|  | *Sporobolus festivus* | 1 |
|  | *Sporobolus fimbriatus* | 10 |
|  | *Sporobolus stapfianus* | 1 |
|  | *Tripogonella minima* | 4 |
|  | *Xerophyta humilis* | 3 |
|  | *Xerophyta retinervis* | 1 |
|  | *Xerophyta viscosa* | 2 |
| CAF | *Microchloa indica* | 1 |
| CHL | *Microchloa indica* | 2 |
|  | *Microchloa kunthii* | 4 |
|  | *Tripogonella spicata* | 5 |
| CHN | *Boea hygrometrica* | 189 |
|  | *Damrongia clarkeana* | 33 |
|  | *Oreocharis mileensis* | 1 |
|  | *Paraboea crassifolia* | 13 |
|  | *Paraboea rufescens* | 115 |
|  | *Microchloa indica* | 13 |
|  | *Microchloa kunthii* | 38 |
|  | *Tripogon filiformis* | 2 |
|  | *Acanthochlamys bracteata* | 39 |
| CIV | *Afrotrilepis pilosa* | 7 |
|  | *Microchloa indica* | 3 |
|  | *Oropetium aristatum* | 1 |
|  | *Sporobolus festivus* | 1 |
|  | *Tripogonella minima* | 1 |
| CMR | *Afrotrilepis pilosa* | 7 |
|  | *Coleochloa abyssinica* | 3 |
|  | *Microdracoides squamosus* | 11 |
|  | *Microchloa indica* | 16 |
|  | *Microchloa kunthii* | 10 |
|  | *Sporobolus festivus* | 7 |
|  | *Tripogon major* | 9 |
|  | *Tripogonella minima* | 10 |
| COD | *Coleochloa abyssinica* | 1 |
|  | *Coleochloa setifera* | 5 |
|  | *Lindernia purpurea* | 1 |
|  | *Microchloa indica* | 11 |
|  | *Microchloa kunthii* | 2 |
|  | *Sporobolus festivus* | 5 |
|  | *Sporobolus stapfianus* | 2 |
| COG | *Microchloa indica* | 1 |
| COL | *Microchloa indica* | 3 |
|  | *Tripogonella spicata* | 1 |
|  | *Vellozia tubiflora* | 21 |
| CUB | *Tripogonella spicata* | 1 |
| ECU | *Microchloa kunthii* | 89 |
|  | *Tripogonella spicata* | 17 |
| ERI | *Craterostigma plantagineum* | 1 |
|  | *Xerophyta schnizleinia* | 1 |
| ESP | *Ramonda myconi* | 50 |
| ETH | *Coleochloa abyssinica* | 10 |
|  | *Craterostigma plantagineum* | 9 |
|  | *Craterostigma pumilum* | 2 |
|  | *Microchloa indica* | 2 |
|  | *Microchloa kunthii* | 4 |
|  | *Sporobolus festivus* | 8 |
|  | *Sporobolus fimbriatus* | 1 |
|  | *Sporobolus pellucidus* | 5 |
|  | *Sporobolus stapfianus* | 1 |
|  | *Tripogonella minima* | 1 |
|  | *Xerophyta humilis* | 1 |
|  | *Xerophyta rippsteinii* | 1 |
|  | *Xerophyta schnizleinia* | 11 |
| FRA | *Ramonda myconi* | 1 |
| GAB | *Afrotrilepis pilosa* | 11 |
| GHA | *Afrotrilepis pilosa* | 2 |
|  | *Microchloa indica* | 7 |
|  | *Microchloa kunthii* | 2 |
|  | *Oropetium aristatum* | 4 |
|  | *Sporobolus festivus* | 1 |
|  | *Tripogonella minima* | 5 |
| GIN | *Afrotrilepis pilosa* | 8 |
|  | *Microchloa indica* | 1 |
| GNB | *Microchloa indica* | 2 |
|  | *Oropetium aristatum* | 1 |
| GNQ | *Afrotrilepis pilosa* | 2 |
| GRC | *Haberlea rhodopensis* | 4 |
|  | *Ramonda nathaliae* | 3 |
|  | *Ramonda serbica* | 2 |
| GTM | *Microchloa kunthii* | 2 |
|  | *Tripogonella spicata* | 1 |
| GUY | *Vellozia candida* | 1 |
|  | *Vellozia tubiflora* | 10 |
| HND | *Microchloa kunthii* | 16 |
|  | *Tripogonella spicata* | 3 |
| IDN | *Paraboea rufescens* | 1 |
| IND | *Eragrostiella bifaria* | 4 |
|  | *Oropetium thomaeum* | 4 |
|  | *Tripogon jacquemontii* | 2 |
| KEN | *Craterostigma hirsutum* | 5 |
|  | *Craterostigma plantagineum* | 1 |
|  | *Linderniella pulchella* | 1 |
|  | *Myrothamnus flabellifolius* | 1 |
|  | *Microchloa indica* | 2 |
|  | *Microchloa kunthii* | 7 |
|  | *Oropetium capense* | 1 |
|  | *Sporobolus festivus* | 1 |
|  | *Sporobolus fimbriatus* | 5 |
|  | *Sporobolus pellucidus* | 5 |
|  | *Sporobolus stapfianus* | 8 |
|  | *Tripogon curvatus* | 4 |
|  | *Xerophyta schnizleinia* | 1 |
|  | *Xerophyta spekei* | 3 |
| LBR | *Afrotrilepis pilosa* | 12 |
| LKA | *Eragrostiella bifaria* | 2 |
|  | *Eragrostiella brachyphylla* | 1 |
|  | *Oropetium thomaeum* | 2 |
| LSO | *Microchloa caffra* | 3 |
|  | *Oropetium capense* | 2 |
|  | *Sporobolus fimbriatus* | 4 |
|  | *Xerophyta viscosa* | 2 |
| MDG | *Coleochloa setifera* | 75 |
|  | *Myrothamnus moschatus* | 124 |
|  | *Microchloa kunthii* | 7 |
|  | *Sporobolus festivus* | 10 |
|  | *Styppeiochloa hitchcockii* | 60 |
|  | *Tripogonella minima* | 1 |
|  | *Xerophyta dasylirioides* | 166 |
|  | *Xerophyta eglandulosa* | 7 |
|  | *Xerophyta nandrasanae* | 5 |
|  | *Xerophyta pinifolia* | 13 |
|  | *Xerophyta pectinata* | 99 |
| MEX | *Microchloa indica* | 3 |
|  | *Microchloa kunthii* | 206 |
|  | *Sporobolus atrovirens* | 31 |
|  | *Tripogonella spicata* | 9 |
| MKD | *Ramonda nathaliae* | 2 |
|  | *Ramonda serbica* | 1 |
| MLI | *Microchloa indica* | 4 |
|  | *Oropetium aristatum* | 4 |
|  | *Oropetium capense* | 1 |
|  | *Tripogonella minima* | 6 |
| MOZ | *Coleochloa pallidior* | 3 |
|  | *Coleochloa setifera* | 7 |
|  | *Linderniella pulchella* | 5 |
|  | *Linderniella wilmsii* | 1 |
|  | *Myrothamnus flabellifolius* | 9 |
|  | *Eragrostis nindensis* | 2 |
|  | *Microchloa kunthii* | 2 |
|  | *Sporobolus festivus* | 4 |
|  | *Sporobolus fimbriatus* | 19 |
|  | *Sporobolus stapfianus* | 5 |
|  | *Tripogonella minima* | 2 |
|  | *Xerophyta humilis* | 1 |
|  | *Xerophyta scabrida* | 3 |
|  | *Xerophyta splendens* | 2 |
|  | *Xerophyta viscosa* | 1 |
| MWI | *Coleochloa setifera* | 1 |
|  | *Linderniella pulchella* | 2 |
|  | *Myrothamnus flabellifolius* | 4 |
|  | *Eragrostis paradoxa* | 5 |
|  | *Microchloa indica* | 2 |
|  | *Microchloa kunthii* | 3 |
|  | *Sporobolus stapfianus* | 3 |
|  | *Xerophyta splendens* | 18 |
| NER | *Microchloa indica* | 7 |
|  | *Tripogonella minima* | 1 |
| NGA | *Afrotrilepis pilosa* | 15 |
|  | *Coleochloa abyssinica* | 2 |
|  | *Microchloa indica* | 17 |
|  | *Sporobolus festivus* | 12 |
|  | *Tripogon major* | 1 |
|  | *Tripogonella minima* | 5 |
| NIC | *Microchloa kunthii* | 6 |
|  | *Tripogonella spicata* | 1 |
| NPL | *Eragrostiella nardoides* | 1 |
| OMN | *Oropetium capense* | 1 |
| PAN | *Vellozia tubiflora* | 5 |
| PER | *Pitcairnia lanuginosa* | 34 |
|  | *Microchloa kunthii* | 22 |
|  | *Tripogonella spicata* | 11 |
|  | *Barbaceniopsis boliviensis* | 1 |
| PNG | *Tripogonella loliiformis* | 5 |
| PRY | *Microchloa indica* | 19 |
|  | *Tripogonella spicata* | 14 |
| SEN | *Afrotrilepis pilosa* | 1 |
|  | *Microchloa indica* | 25 |
|  | *Oropetium aristatum* | 5 |
|  | *Tripogonella minima* | 4 |
| SLE | *Afrotrilepis pilosa* | 10 |
|  | *Microdracoides squamosus* | 2 |
|  | *Microchloa indica* | 1 |
|  | *Tripogon major* | 4 |
| SOM | *Oropetium thomaeum* | 1 |
|  | *Sporobolus pellucidus* | 1 |
|  | *Sporobolus ruspolianus* | 6 |
|  | *Xerophyta schnizleinia* | 1 |
| SRB | *Ramonda nathaliae* | 1 |
|  | *Ramonda serbica* | 1 |
| SWZ | *Coleochloa setifera* | 1 |
|  | *Linderniella pulchella* | 1 |
|  | *Linderniella wilmsii* | 6 |
|  | *Microchloa kunthii* | 4 |
|  | *Sporobolus stapfianus* | 6 |
| TCD | *Microchloa indica* | 1 |
|  | *Sporobolus festivus* | 1 |
|  | *Tripogonella minima* | 1 |
| TGO | *Afrotrilepis pilosa* | 1 |
|  | *Oropetium aristatum* | 1 |
|  | *Tripogonella minima* | 1 |
| THA | *Paraboea rufescens* | 5 |
|  | *Microchloa indica* | 3 |
| TZA | *Coleochloa abyssinica* | 4 |
|  | *Coleochloa microcephala* | 15 |
|  | *Coleochloa setifera* | 5 |
|  | *Craterostigma hirsutum* | 12 |
|  | *Craterostigma plantagineum* | 5 |
|  | *Craterostigma pumilum* | 4 |
|  | *Linderniella wilmsii* | 2 |
|  | *Myrothamnus flabellifolius* | 7 |
|  | *Eragrostiella bifaria* | 4 |
|  | *Eragrostis nindensis* | 2 |
|  | *Microchloa caffra* | 6 |
|  | *Microchloa indica* | 5 |
|  | *Microchloa kunthii* | 35 |
|  | *Oropetium capense* | 3 |
|  | *Oropetium thomaeum* | 4 |
|  | *Sporobolus festivus* | 37 |
|  | *Sporobolus fimbriatus* | 39 |
|  | *Sporobolus pellucidus* | 15 |
|  | *Sporobolus stapfianus* | 12 |
|  | *Tripogon major* | 8 |
|  | *Tripogonella minima* | 1 |
|  | *Xerophyta scabrida* | 14 |
|  | *Xerophyta spekei* | 10 |
| UGA | *Craterostigma plantagineum* | 4 |
|  | *Microchloa indica* | 1 |
|  | *Microchloa kunthii* | 7 |
|  | *Sporobolus festivus* | 11 |
|  | *Sporobolus fimbriatus* | 1 |
|  | *Sporobolus stapfianus* | 5 |
| USA | *Microchloa kunthii* | 1 |
|  | *Tripogonella spicata* | 4 |
| VEN | *Trilepis lhotzkiana* | 1 |
|  | *Microchloa indica* | 2 |
|  | *Tripogonella spicata* | 3 |
|  | *Vellozia tubiflora* | 83 |
| VNM | *Paraboea crassifolia* | 8 |
|  | *Paraboea rufescens* | 7 |
|  | *Microchloa indica* | 1 |
| ZAF | *Coleochloa setifera* | 59 |
|  | *Linderniella pulchella* | 9 |
|  | *Linderniella wilmsii* | 37 |
|  | *Eragrostis nindensis* | 10 |
|  | *Microchloa caffra* | 68 |
|  | *Microchloa kunthii* | 97 |
|  | *Oropetium capense* | 156 |
|  | *Sporobolus festivus* | 2 |
|  | *Sporobolus fimbriatus* | 125 |
|  | *Sporobolus stapfianus* | 113 |
|  | *Tripogonella minima* | 38 |
|  | *Xerophyta elegans* | 63 |
|  | *Xerophyta retinervis* | 4 |
|  | *Xerophyta schlechteri* | 1 |
|  | *Xerophyta villosa* | 11 |
|  | *Xerophyta viscosa* | 1 |
| ZMB | *Coleochloa setifera* | 2 |
|  | *Linderniella wilmsii* | 1 |
|  | *Myrothamnus flabellifolius* | 1 |
|  | *Eragrostis nindensis* | 3 |
|  | *Eragrostis paradoxa* | 2 |
|  | *Microchloa caffra* | 2 |
|  | *Microchloa indica* | 9 |
|  | *Microchloa kunthii* | 6 |
|  | *Micrachne patentiflora* | 2 |
|  | *Sporobolus festivus* | 1 |
|  | *Sporobolus fimbriatus* | 2 |
|  | *Sporobolus stapfianus* | 2 |
|  | *Xerophyta villosa* | 1 |
| ZWE | *Coleochloa pallidior* | 1 |
|  | *Coleochloa setifera* | 5 |
|  | *Linderniella pulchella* | 9 |
|  | *Eragrostis nindensis* | 3 |
|  | *Eragrostis paradoxa* | 2 |
|  | *Microchloa indica* | 4 |
|  | *Microchloa kunthii* | 24 |
|  | *Micrachne patentiflora* | 12 |
|  | *Sporobolus festivus* | 1 |
|  | *Sporobolus fimbriatus* | 8 |
|  | *Sporobolus stapfianus* | 24 |
|  | *Xerophyta humilis* | 3 |
|  | *Xerophyta schlechteri* | 1 |

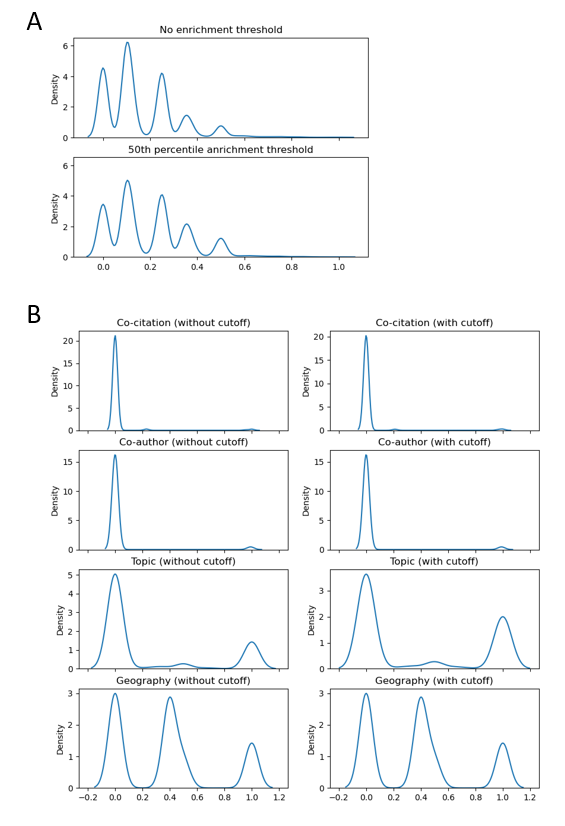

**Supplemental Figure 1. Score distributions with and without applying the 50th percentile enrichment cutoff.** Distributions of (a) composite and (b) single scores.

**Supplemental note 1. Algorithm 1 Basic conference invitee recommendation algorithm**

**Algorithm 1** Basic conference invitee recommendation algorithm

*>* Build networks and topic model

*co-citation network*

*co-author network*

*topic clusters*

> Cluster networks hierarchically

*co-citation clusters* ← Louvain(*co-citation network*)

*co-author clusters* ← Louvain(*co-author network*)

> Calculate the mean distance between each author and all clusters

**for** *cluster type* in *[co-citation clusters, co-author clusters]* **do**

**for** *author* in *dataset* **do**

**for** *cluster* in *cluster type* **do**

*distance ←* len(*path to cluster*) + len(*path to author*) - 2*(*common path length*)

**end for**

*mean distance ←* mean(*distances*)

**end for**

**end for**

*>* Calculate cluster enrichments

**for** *cluster type* in *[co-citation clusters, co-author clusters, topic clusters]* **do**

**for** *cluster* in *cluster type* **do**

*cluster enrichment* ← *num conference authors in cluster* / *num authors in cluster*

**end for**

**end for**

> Calculate geographic conference enrichment

**for** *country* in *world* **do**

*geographic conference enrichment ← number attendees from country* / *number attendees*

*top 50p countries ←* percentile(*nonzero* *geographic conference enrichments*, 50)

*bottom 50p countries ← country* **for** *country* in *nonzero countries* **if** *country* not in *top 50p countries*

> Apply calculation of candidate scores

**for** *author* in *dataset* **do**

> Calculate score

*all scores ←* calculate scores(*cluster distances, enrichments*)

> Average all scores

**if** *author* not in *previous attendees* **then**

*author overall score ←* mean(*all scores*)

**else**

*author overall score ←* 0

**end if**

**end for**

> Rank and apply cutoff

*sorted authors* ← sort(*author overall scores*)

*candidates ←* top X percent(*sorted authors*)

**return** *candidates*

**Supplemental note 2. Algorithm 1 Basic conference invitee recommendation algorithm**

**Algorithm 2** Candidate score calculation

> Hierarchical cluster scores

**for** *distances* in *[co-citation distances, co-author distances]* **do**

*penalty threshold ←* percentile(*distances*, 25)

*close side distance ← penalty threshold - 0*

*far side distance ←* max(*distances*) - *penalty threshold*

**for** *distance* in *distances* **do**

*threshold distance ← penalty threshold - distance*

**if** *threshold distance* < 0 **then**

*normalized distance ←* 1 - (abs(*threshold distance*)/*far side distance*

**else**

*normalized distance ←* 1 - (*threshold distance*/*close side distance*)

**end if**

**end for**

*mean cluster score ←* mean(*normalized distances*)

**end for**

> Topic model scores (optimized for topic novelty)

**for** *topic* in *topic clusters* **do**

*topic score ←* 1 **if** *topic* not in *enriched topics* **else** 0

**end for**

*mean topic score ←* mean(*topic scores*)

> Geographic score (optimized for geographic novelty)

**if** *author country* in *top 50p countries* **then**

*geographic score ←* 0

**else if** *author country* in *bottom 50p countries* **then**

*geographic score ←* 0.5

**else if** *author* does not have *country* **then**

*geographic score ← general probability score*

**else**

*geographic score ←* 1

**end if**

**return** *all scores*
